# Combinatorial chloride and calcium channelopathy in myotonic dystrophy

**DOI:** 10.1101/2023.05.29.542752

**Authors:** Lily A. Cisco, Matthew T. Sipple, Katherine M. Edwards, Charles A. Thornton, John D. Lueck

## Abstract

Myotonic dystrophy type 1 (DM1) involves misregulated alternative splicing for specific genes. We used exon or nucleotide deletion to mimic altered splicing of genes central to muscle excitation-contraction coupling processes in mice. Mice with forced-skipping of exon 29 in Ca_V_1.1 calcium channel combined with loss of ClC-1 chloride channel function showed a markedly reduced lifespan, whereas other combinations of splicing mimics did not affect survival. The Ca^2+^/Cl^-^ bi-channelopathy mice exhibited myotonia, weakness, and impairment of mobility and respiration. Chronic administration of the calcium channel blocker verapamil rescued survival and improved force generation, myotonia, and respiratory function. These results suggest that Ca^2+^/Cl^-^ bi-channelopathy contributes to muscle impairment in DM1 and is potentially mitigated by common clinically available calcium channel blockers.

**Summary:** Repurposing of a calcium channel blocker extends life and mitigates muscle and respiratory dysfunction in a myotonic dystrophy type 1 Ca^2+^/Cl^-^ bi-channelopathy mouse model.

## Introduction

Myotonic dystrophy type 1 (DM1) is an autosomal dominant disorder characterized by myotonia, progressive weakness, heart block, gastrointestinal dysmotility, predisposition to malignancy, and other symptoms(*1*). Survival is limited by weakness of the respiratory muscles or cardiac arrhythmia. The genetic basis is an expansion of CTG repeats in the 3′ untranslated region of the *DM1 Protein Kinase* (*DMPK*) gene. Although DM1 can begin in infancy, more often the symptoms are delayed to the 2^nd^-5^th^ decade. The onset and progression of DM1 is believed to reflect an age-dependent increase in the size of expanded repeats in tissue, reaching lengths of several thousand repeats in skeletal and cardiac muscle.

The core mechanism for DM1 is RNA toxicity, whereby *DMPK* transcripts with expanded CUG repeats form nuclear condensates that sequester splicing factors in the Muscleblind-like (MBNL) family. The resulting loss of MBNL function causes misregulation of alternative splicing and other changes of RNA processing. This in turn produces a complex array of clinical findings in which specific endophenotypes such as myotonia, heart block, or insulin resistance, are linked to splicing defects of particular genes, such as ClC-1 chloride channel(*2-4*), SCN5A sodium channel(*5, 6*), or insulin receptor(*7*).

Efforts to deconstruct the DM1 phenotype have taken a reductionist approach in which individual splicing defects are studied in isolation(*8, 9*). However, the main source of disability – progressive muscle weakness – remains largely unexplained. It is unclear whether motor disability results from predominant effects of an individual splicing defect or combinatorial mis-splicing of several or many transcripts. Although splicing misregulation of many alternative exons correlates with muscle weakness, it has been difficult to establish causal relationships because the splicing changes are highly intercorrelated(*10*). Accordingly, we used exon or nucleotide deletion to mimic splicing defects affecting four genes involved in sequential steps of excitation contraction coupling (ECC), including *Clcn1* (encoding ClC-1 chloride channel), *Cacna1s* (Ca_V_1.1 calcium channel, dihydropyridine receptor), *Ryr1* (ryanodine receptor, Ca^2+^ release channel), and *Atp2a1* (sarcoplasmic reticulum Ca^2+^-ATPase, Serca1, Ca^2+^ reuptake pump). Then we used breeding to examine combinatorial effects. We found that ClC-1/Ca_V_1.1 bi-channelopathy, but not other combinations, was highly deleterious, causing weakness, respiratory impairment, and marked reduction of lifespan. The survival of bi-channelopathy mice was dramatically improved by chronic oral feeding of the L-type calcium channel blocker, verapamil. These results support a contributory role for excess calcium entry in DM1 pathogenesis and suggests that Ca^2+^ channel modulation is a potential therapeutic strategy.

## Results

### Generation of forced spliced DM1 mouse models

ECC stands out among pathways affected by DM1 mis-splicing because three sequential components are strongly affected by exon skipping (*CACNA1S* exon 29, *RYR1* exon 70, and *ATP2a1* exon 22) and a fourth shows abnormal exon inclusion (*CLCN1* exon 7a). To reproduce each effect we generated congenic lines having genomic deletion of the DM1-skipped exons, RyR1^Δe70^, Serca1a^Δe22^, or Ca_V_1.1^Δe29^, and then verified production of the expected splice products by cDNA sequencing. To reproduce effects of *CLCN1* mis-splicing we used a single nucleotide deletion (*adr*^mto2j^ mice) that, like exon 7a inclusion, causes frameshift, protein truncation, and loss of channel function (ClC-1^-/-^)(*11*). We found that all splicing mimics studied in isolation, whether in the heterozygous or homozygous state, exhibited normal cage behavior, weight gain, and survival, except that homozygous ClC-1^-/-^ mice showed the expected phenotype of muscle stiffness (myotonia) and modest reduction of body weight, with average survival of ∼35 weeks in our colony.

Next we tested for combinatorial effects. Through breeding we obtained mice doubly homozygous for each possible combination of two splicing mimics (*n* = 7 combinations), and for the triple combination of RyR1^Δe70^, Serca1a^Δe22^, and Ca_V_1.1^Δe29^. One combination emerged as highly deleterious. The mean lifespan of Ca_V_1.1^Δe29/Δe29^/ClC-1^-/-^ double homozygotes was 8.1 weeks (Fig. 1A). Even with heterozygosity for Ca_V_1.1^Δe29^, the mean lifespan on the ClC-1^-/-^ background was 9.8 weeks, with no mice surviving past 14 weeks. As other combinations did not show reduced survival or failure to gain weight that differed from ClC-1^-/-^ mice (Fig. 1B,C), our subsequent experiments focused on the Ca^2+^/Cl^-^ bi-channelopathy mice.

**Fig. 1.**
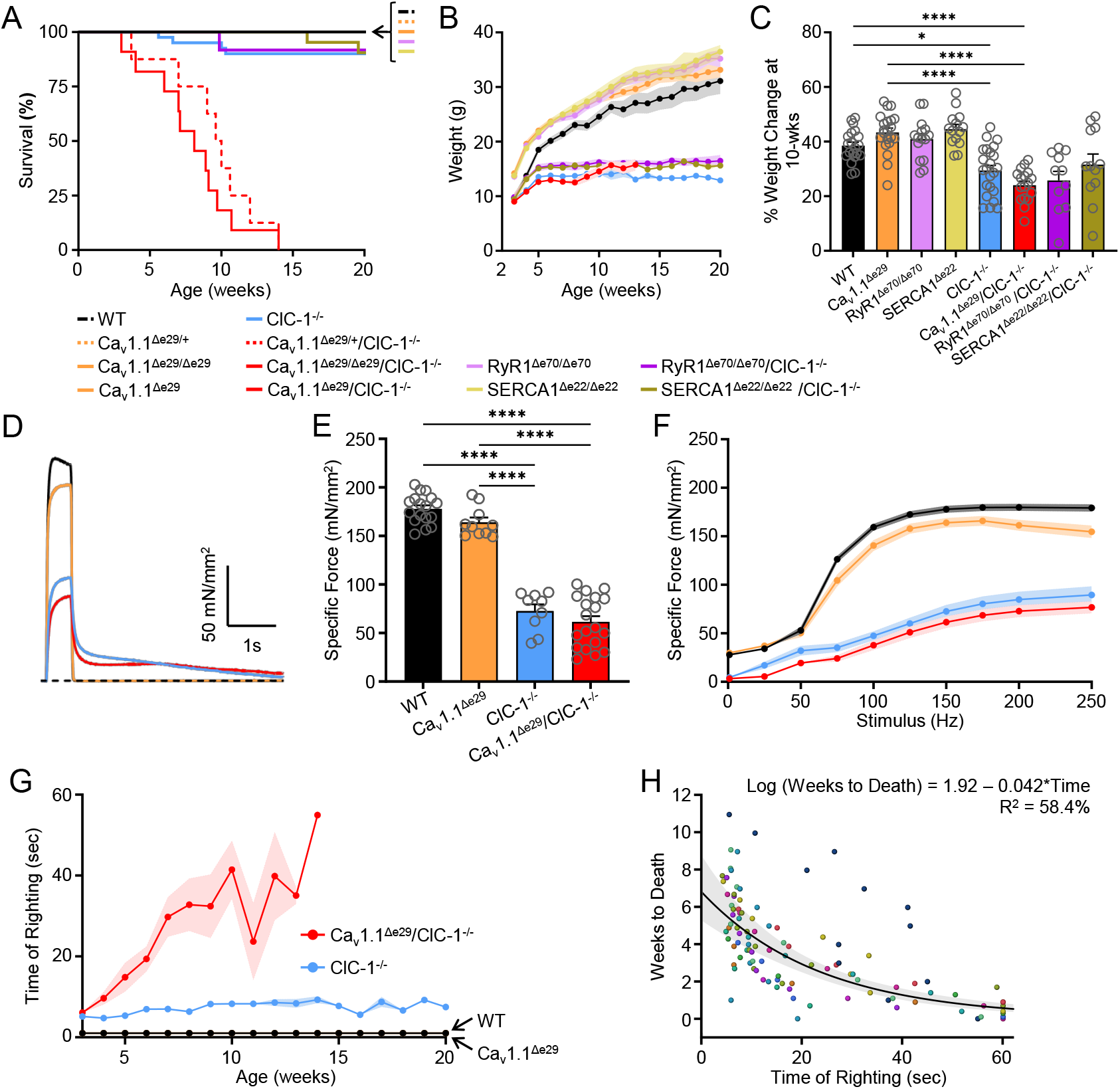
Ca_V_1.1^Δe29^ and ClC-1^-/-^ alleles exhibit synthetic lethality, and result in significantly reduced body weight and severe muscle weakness in mice. **A)** Kaplan-Meier survival analysis of WT (n=10; female=5, male=5), Ca_v_1.1^Δe29/+^ (n=13; female=7, male=6), Ca_v_1.1^Δe29^ (n=15; female=7, male=8), RyR1^Δe70/Δe70^ (n=21; female=10, male=11), SERCA1^Δe22/Δe22^ (n=19; female=10, male=9), ClC-1^-/-^ (n=40; female=21, male=19), RyR1^Δe70/Δe70^/ClC-1^-/-^ (n=12; female=6, male=6), SERCA1^Δe22/Δe22^/ClC-1^-/-^ (n=22; female=11, male=11), Ca_v_1.1^Δe29/+^/ClC-1^-/-^ (n=8; female=3, male=5), and Ca_v_1.1^Δe29/Δe29^/ClC-1^-/-^ (n=11; female=6, male=5). Of note, non-ClC-1^-/-^ mouse combinations are not shown **B)** Weekly body weight analysis. **c)** Percent body weight change from weaning at 10 wks. **B, C)** WT (n=20; female=10, male=10), Ca_v_1.1^Δe29^ (n=21; female=12, male=9), RyR1^Δe70/Δe70^ (n=15; female=5, male=10), SERCA1^Δe22/Δe22^ (n=14; female=7, male=7), ClC-1^-/-^ (n=23; female=13, male=10), Ca_v_1.1^Δe29^/ClC-1^-/-^ (n=17; female=8, male=9), RyR1^Δe70/Δe70^/ClC-1^-/-^ (n=11; female=5, male=6), SERCA1^Δe22/Δe22^/ClC-1^-/-^ (n=12; female=8, male=4). **D)** Representative specific force traces and **E)** average peak specific force elicited by 150Hz (500ms) tetanic stimulation of isolated EDL muscles from 10-wk mice. **F)** Average frequency dependence of specific force generation, elicited from isolated EDL muscle from 10-wk mice. **E, F)** WT (n=17; female=8, male=9), Ca_v_1.1^Δe29^ (n=10; female=5, male=5), ClC-1^-/-^ (n=9; female=4, male=5), Ca_v_1.1^Δe29^/ClC-1^-/-^ (n=19; female=8, male=11). **G)** Weekly time of righting reflex analysis of Ca_v_1.1^Δe29^/ClC-1^-/-^ (red), ClC-1^-/-^ (light blue), Ca_v_1.1^Δe29^ (orange), WT (black). **H)** Correlation of time of righting reflex to weeks till death of Ca_v_1.1^Δe29^/ClC-1^-/-^ mice (n=15; female=7, male=8). Different colors of the dots represent individual mice, repeats of the same color are individual time of righting measures. Symbols, open circles, individual mice; bars and closed circles, means ± SEM. Statistical analysis of results in Fig. 1. are found in Supplementary Notes.

These results indicated that exon 29 skipping for Ca_V_1.1 is highly deleterious in the context of ClC-1 loss and suggested that Ca_V_1.1^Δe29^ channels exert a dominant gain-of-function. To test the latter possibility we used whole-cell patch clamp to isolate macroscopic Ca_V_1.1 currents in *flexor digitorum brevis* (FDB) muscle fibers. As compared to WT littermates, Ca_V_1.1 channel activity in homozygous Ca_V_1.1^Δe29/Δe29^ fibers exhibited a significant left-shift in activation (∼25mV) and an augmented peak current-density (Supplementary Fig. S1A,B,C), comparable to previous findings in homozygous Ca_V_1.1^Δe29/Δe29^ mice that were also obtained by exon deletion and in heterologous expression studies.(*9, 12*) Surprisingly, we also found that abnormalities of Ca_V_1.1 channel activity in heterozygous Ca_V_1.1^Δe29/+^ fibers were indistinguishable from homozygotes, confirming a dominant gain-of-function by Ca_V_1.1^Δe29^ channels. By RT-PCR analysis we found that dominant behavior of Ca_V_1.1^Δe29^ channels in Ca_V_1.1^Δe29/+^ heterozygous mice did not result from unequal allelic accumulation of Ca_V_1.1 mRNA (Supplementary Fig. S1D). Taken together, these results suggest that increased muscle activity (myotonic runs due to ClC-1 loss) plus increased calcium entry during muscle activity (via Ca_V_1.1^Δe29^ channels) causes premature death in bi-channelopathy mice.

Next, we examined the effects of single vs. dual channelopathy on *ex vivo* force generation in hindlimb muscle. Since Ca_V_1.1 activity was similar in Ca_V_1.1^Δe29/+^ and Ca_V_1.1^Δe29/Δe29^ mice, these genotypes were grouped together (“Ca_V_1.1^Δe29^”) for these and subsequent analyses. Single-twitch and tetanic stimulation of *extensor digitorum longus* (EDL) muscle showed similar reduction of maximum force in single- (ClC-1^-/-^, Fig. 1D-F, blue) and bi-channelopathy (Ca_V_1.1^Δe29^/ClC-1^-/-^, Fig. 1D-F, red) muscle, as compared to WT (Fig. 1D-F, black) and Ca_V_1.1^1e29^ (Fig. 1D-F, orange) controls. Representative traces of tetanic force (150Hz, Fig. 1D) also demonstrated myotonia, evidenced by delayed muscle relaxation in ClC-1^-/-^ (blue trace) and Ca_V_1.1^Δe29^/ClC-1^-/-^ (red trace) muscle, which was absent in WT (black trace) and Ca_V_1.1^Δe29^ (orange trace) controls.

These results under conditions of *ex vivo* tetanic stimulation suggested fixed weakness in ClC-1 null muscle, as observed with age in patients with recessive myotonia congenita due to biallelic mutations of ClC-1(*13*), but showed no difference of contractility to account for premature mortality in bi- vs. single-channelopathy mice. Muscle histochemistry and immunofluorescence at 10 weeks showed the expected adaptive response to myotonia in ClC^-/-^ muscles, such as fiber type switching and increased oxidative activity, but also failed to distinguish single-from bi-channelopathy mice. Further, the bi-channelopathy mice did not show a flagrant myopathy or histologic features that are characteristic of human DM1, such as, central nuclei or circumferential myofibrils (ring fibers) (Supplementary Fig. S2-6). However, the mobility of bi-channelopathy mice was strikingly affected. A well-known feature of ClC-1 null mice, also known as *arrested development of righting response* (*adr*) mice, is the slow recovery of upright posture after being placed supine(*14-16*), and this righting response was clearly delayed in ClC-1^-/-^ mice compared to WT or Ca_V_1.1^Δe29^ controls (Fig. 1G, blue). However, the time of righting response (TRR) was much greater in bi-channelopathy mice, and this deficit was progressive (Fig. 1G, red). Indeed, a rise of righting response beyond 30 sec was predictive of death or survival endpoint in bi-channelopathy mice (Fig. 1H).

### Ca_V_1.1^Δe29^ conductance potentiates muscle transient weakness in context of ClC-1 loss

The results above indicated that ClC-1^-/-^ mice, regardless of Ca_V_1.1 splicing, show weakness of hindlimb muscles during brief (0.5 s) tetanic stimulation *ex vivo*, designated here as ‘fixed weakness’. Previous studies of recessive myotonia in humans and mice have also shown a superimposed transient weakness. This reversible weakness occurs when fibers with genetic loss or pharmacologic block of ClC-1 are rendered temporarily inexcitable (depolarized) by sustained or repetitive muscle activity.(*17-20*) In WT fibers under conditions of ClC-1 blockade, the transient weakness was shown to depend on Ca^2+^ currents through normal (e29^+^) Ca_V_1.1 channels(*20*). We therefore postulated that e29 skipping and resulting enhanced Ca^2+^ current may further aggravate transient weakness. We tested this possibility using *ex vivo* stimulation of EDL. Muscles from 6-week old ClC-1^-/-^ and Ca_V_1.1^Δe29^/ClC-1^-/-^ mice were subjected to a series of brief stimuli (0.5 s × 100 Hz) at 4 s intervals for 3 minutes (Fig. 2). Representative ClC-1^-/-^ (Fig. 2A, blue trace) and Ca_V_1.1^Δe29^/ClC-1^-/-^ (Fig. 2A, red trace) force traces of the first 15 stimulations, normalized to maximum force of the first stimulation, demonstrated a transient drop of force production after the first stimulation, which was followed by recovery over subsequent stimulations. The transient weakness in Ca_V_1.1^Δe29^/ClC-1^-/-^ EDLs exhibited a ∼75% drop in force (Fig 2B red symbols), compared to ∼60% in ClC-1^-/-^ EDLs (Fig 2B, blue symbols).

**Fig. 2.**
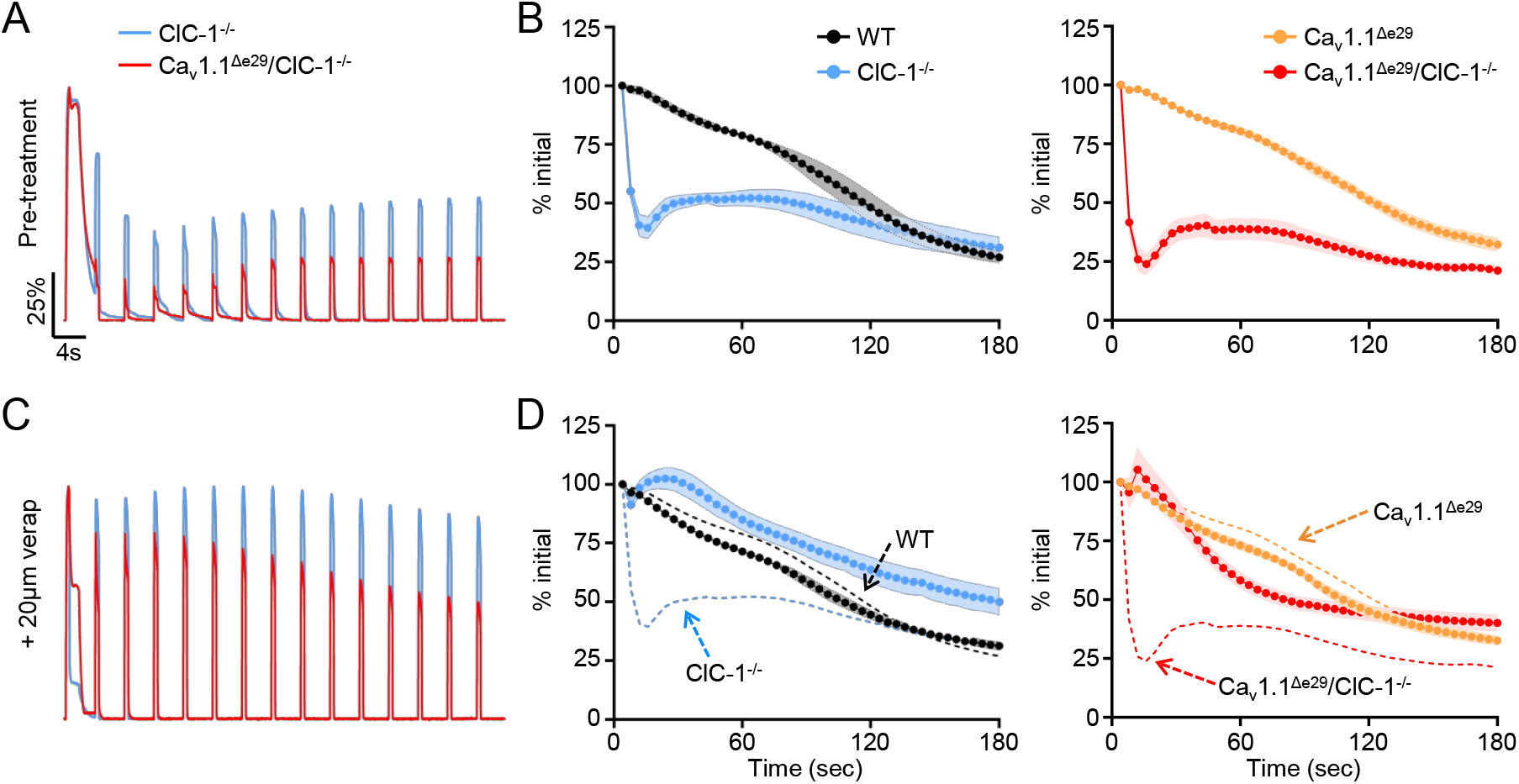
Ca_V_1.1^Δe29^/ClC-1^-/-^ and ClC-1^-/-^ muscle exhibits severe transient weakness that is significantly improved by the addition of verapamil. **A, C)** Normalized representative force traces of the first 15 tetani (100Hz, 500ms) separated by four seconds, recorded *ex vivo* from EDLs isolated from 6-wk ClC-1^-/-^ (blue) and Ca_V_1.1^Δe29^/ClC-1^-/-^ (red) mice in the **A)** absence and **C)** presence of 20mM verapamil added to the bath. **B, D)** Average peak tetanic EDL forces normalized to the initial stimulus, elicited by 44 subsequent 100Hz, 500ms tetanic stimulations separated by four seconds from 6-wk WT (black, n=4), ClC-1^-/-^ (blue n=4), Ca_V_1.1^Δe29^ (orange, n=4) and Ca_V_1.1^Δe29^/ClC-1^-/-^ red, n=4) mice in the **B)** absence and **D)** presence of 20mM verapamil added to the bath for ClC-1^-/-^ (blue n=4) and Ca_V_1.1^Δe29^/ClC-1^-/-^ red, n=4) EDLs. Dashed lines in **D)** represent average data presented in **B)** as a reference for pre-treatment. **B, D)** WT (n=4; female=2, male=2), Ca_v_1.1^Δe29^ (n=4; female=2, male=2), ClC-1^-/-^ (n=4; female=2, male=2), Ca_V_1.1^Δe29^/ClC-1^-/-^ (n=4; female=2, male=2). Symbols, closed circles, mean ± SEM. Statistical analysis of results in Figure 2. are found in Supplementary Notes.

Next, we wanted to determine if exaggerated transient weakness in Ca_V_1.1^Δe29^/ClC-1^-/-^ muscle depends on Ca^2+^ entry and responds to pharmacologic block of Ca_V_1.1. Verapamil is a phenylalkylamine compound in wide clinical use for blocking Ca_V_1.2 channels in cardiac and smooth muscle(*21-24*). It is also known to block Ca_V_1.1 channels in skeletal muscle(*24*) but this is not considered clinically relevant because the Ca^2+^ conductance of Ca_V_1.1 is dispensable for ECC of normal skeletal muscle(*25*). Remarkably, addition of 20 μM verapamil to the bath completely abrogated the transient weakness in ClC-1^-/-^ and Ca_V_1.1^Δe29^/ClC-1^-/-^ EDL muscles *ex vivo* (Fig 2C,D, blue and red, respectively). In contrast, addition of verapamil to the bath did not recover the fixed component of weakness in ClC-1^-/-^ or Ca_V_1.1^Δe29^/ClC-1^-/-^ muscles, as shown by non-normalized force plots in Supplementary Fig. S7C,D. Next, we determined the impact of Ca_V_1.1^Δe29^ conductance on transient weakness in the setting of pharmacologic inhibition of ClC-1, where the long-term adaptive response of skeletal muscle to myotonia has not occurred(*26*). We blocked ClC-1 chloride channels in WT and Ca_V_1.1^Δe29^ EDL muscles with 100 μM 9-anthracene carboxylic acid (9-AC), which is known to block more than 95% of ClC-1 channels in mouse skeletal muscle(*27*). Notably, this concentration of 9-AC only produced slight transient weakness in WT muscle (Supplementary Fig. S8A,B, blue trace and symbols), which likely reflects the incomplete block of Cl^-^ conductance. In contrast, application of 9-AC to Ca_V_1.1^Δe29^ muscle produced severe transient weakness, eliminating 95% of force production, which then recovered to 55% of the original force after 60 seconds before exhibiting the expected muscle fatigue (Supplementary Fig. S8A,B, red trace and symbols). Once again, the addition of verapamil strongly inhibited transient weakness in a dose dependent manner (Supplementary Fig. S8C-F). Taken together with previous work(*20*), these results confirm that Ca^2+^ currents through Ca_V_1.1 underlie transient weakness in ClC-1 null muscles, and show that this effect is enhanced for Ca_V_1.1^Δe29^ channels and rescued by Ca^2+^ channel block.

### Ca_V_1.1^Δe29^ conductance potentiates chloride channel myotonia

Given the impact of Ca_V_1.1^Δe29^ channels on transient weakness and the interplay of transient weakness and myotonia, we also wanted to determine the effects of Ca_V_1.1^Δe29^ on myotonia in ClC-1 deficient fibers. Using EDL muscles isolated from 6-week old mice, myotonia was taken as the inappropriate continuation of force production after tetanic stimulation was stopped (Fig. 3). We elicited muscle contraction using three successive rounds of tetanic stimulation (150Hz, 500ms), each separated by 3 minutes of rest. This rest period reduces the well-known ‘warm-up’ phenomenon of chloride channel myotonia(*15*), in which myotonia wanes with repeated bouts of muscle activation. The force traces were normalized to peak force and then integrated over time to quantify myotonia (dashed lines in Fig 3A,C,G,E). The integrated signals for 1^st^, 2^nd^ and 3^rd^ stimulations were then compared (Fig. 3B,D,F,H). As expected, myotonia was not observed in WT (black) or Ca_V_1.1^Δe29^ (orange) muscles (Fig. 3A,B). In contrast, ClC-1^-/-^ (blue) and Ca_V_1.1^Δe29^/ ClC-1^-/-^ (red) muscles both showed robust myotonia, but severity was greater in bi-channelopathy muscle (Fig. 3E,F). Interestingly, despite the 3-minute rest interval, the myotonia in Ca_V_1.1^Δe29^/ClC-1^-/-^ but not in ClC-1^-/-^ muscle demonstrated ‘warm-up’, showing a 42% decrease over successive stimulations (Fig. 3F, red) without any corresponding drop in peak specific force (Supplementary Fig S9). Contralateral muscles were examined in the presence of 20 μM verapamil. Remarkably, the myotonic after-contractions were virtually eliminated by verapamil in both ClC-1^-/-^ (blue) and Ca_V_1.1^Δe29^/ClC-1^-/-^ (red) EDL (Fig. 3G,H and Supplementary Fig. S10B). We also compared the impact of Ca_V_1.1^Δe29^ channels on myotonia brought about by acute pharmacologic blockade of ClC-1 channels (Supplementary Fig. S11). As with our studies of transient weakness, we observed differences between genetic and pharmacologic myotonia in both WT and Ca_V_1.1^Δe29^ muscle. Comparing the integrated normalized force production, the severity of myotonia is significantly less in 9-AC treated WT and Ca_V_1.1^Δe29^ muscles (Fig. 3F) as compared to genetic counterparts (ClC-1^-/-^ and Ca_V_1.1^Δe29^/ClC-1^-/-^) (Supplementary Fig. S12D; comparative analysis in Supplementary Fig. S12). Unlike Ca_V_1.1^Δe29^/ClC-1^-/-^ muscle, there was no indication of ‘warm-up’ in 9-AC treated Ca_V_1.1^Δe29^ muscle from the 1^st^ to 3^rd^ stimulus series (Supplementary Fig. S11D). Further, there is no observed myotonia ‘rebound’ observed in 9-AC treated WT and Ca_V_1.1^Δe29^ EDLs (Supplementary Fig. S11D), where it is observed in all force recordings from ClC-1^-/-^ and Ca_V_1.1^Δe29^/ClC-1^-/-^ EDLs (Supplementary Fig. S10A). Regardless of these differences, application of 5 and 20 μM verapamil again showed dose-dependent reduction of 9-AC-induced myotonia in both WT and Ca_V_1.1^Δe29^ EDLs, (Supplementary Fig. S11E,F), without a significant impact on peak force production (Supplementary Fig. S13). Taken together, these results indicate that Ca^2+^ conductance potentiates myotonia in the context of normal Ca_V_1.1^+e29^ channels. This effect is further aggravated when Ca_V_1.1 exon 29 is skipped, and in both circumstances the myotonia is mitigated by verapamil. It is also apparent that Ca_V_1.1^Δe29^ unmasks myotonia even when loss of ClC-1 conductance (pharmacologic block) is incomplete, and that increased Ca^2+^ conductance through Ca_V_1.1^Δe29^ appears to enhance the warmup phenomenon.

**Fig. 3:**
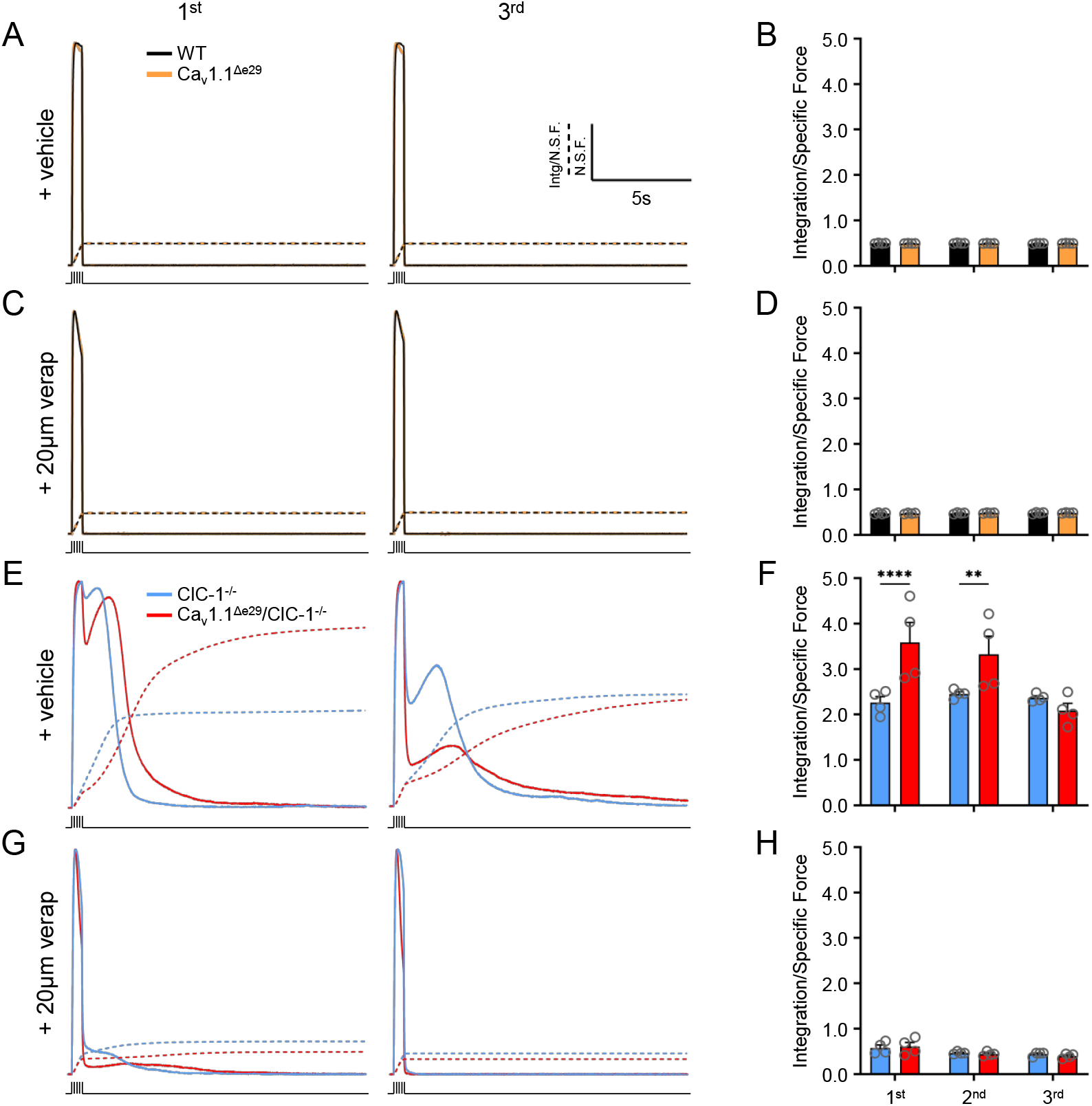
Verapamil significantly reduces myotonia in both Ca_v_1.1^Δe29^/ClC-1^-/-^ and ClC-1^-/-^ mouse muscle. **A, C)** Normalized representative specific force traces of the first (left) and third (right) tetani (150Hz, 500ms) from EDLs isolated from 6-wk WT (black) and Ca_v_1.1^Δe29^ (orange) mice in the **A)** absence and **C)** presence of 20mM verapamil added to the bath. Dashed lines represent accumulated force. **B, D)** Average normalized integration of force for WT (black) and Ca_v_1.1^Δe29^ (orange) EDLs across 3 tetanic stimulations (150Hz, 500ms) in the **B)** absence and **D)** presence of 20 mM verapamil added to the bath. **E, G)** Normalized representative specific force traces of the first (left) and third (right) tetani (150Hz, 500ms) from EDLs isolated from 6-wk ClC-1^-/-^ (blue) and Ca_v_1.1^Δe29^/ClC-1^-/-^ (red) mice in the absence **E)** and presence **G)** of 20 mM verapamil added to the bath. Dashed lines represent accumulated force. **F, H)** Average integration of force for ClC-1^-/-^ (blue) and Ca_v_1.1^Δe29^/ClC-1^-/-^ (red) EDLs across 3 tetanic stimulations (150Hz, 500ms) in the **B)** absence and **D)** presence of 20 mM verapamil added to the bath. **B, D, F, H)** WT (n=4; female=2, male=2), Ca_v_1.1^Δe29^ (n=4; female=2, male=2), ClC-1^-/-^ (n=4; female=2, male=2), Ca_v_1.1^Δe29^/ClC-1^-/-^ (n=4; female=2, male=2). Symbols, open circles, individual mice; bars, mean ± SEM. Statistical analysis of results in Fig. 3. are found in Supplementary Notes. *Notes: Contralateral EDLs were used for each untreated and treated experiment. All traces plotted with the same scale of time and normalized force for comparison. The same traces without force normalization and with expanded timescales for closer examination of myotonia are shown in Supplemental Figures 9 and 10, respectively*.

### Long-term verapamil administration improves survival, body weight and motility of Ca_V_1.1^Δe29^/ClC-1^-/-^ mice

Since short-term exposure to verapamil inhibited transient weakness and myotonia *ex vivo* in Ca_V_1.1^Δe29^/ClC-1^-/-^ muscle, we wanted to determine if long-term administration *in vivo* would improve survival and muscle function in bi-channelopathy mice. Previous work has shown that oral administration of verapamil mixed in food at an estimated dose of 100 mg/kg/d was well tolerated over 16 weeks in mice lacking the sarcoglycan-sarcospan complex in smooth muscle, and rescued the cardiomyopathy caused by microvascular defects(*28*). In a pilot study of oral feeding we found that estimated regimens of 100 and 200 mg/kg/d of verapamil, starting at the time of weaning (3-weeks of age), were well tolerated in Ca_V_1.1^Δe29^/ClC-1^-/-^ mice, and appeared to extend survival (*n* = 3 or 4, respectively, Supplementary Fig. S14). Since 3 of 4 mice on 200mg/kg/d survived to 20 weeks we carried this regimen forward in a repeat study of a larger cohort. Once again, verapamil significantly extended survival in Ca_V_1.1^Δe29^/ClC-1^-/-^ mice. 8 of 9 mice survived beyond 14 weeks, which was never observed in untreated mice receiving the same diet, and 7 of 9 survived to the age of 20 weeks (Fig. 4A, green line) when they were euthanized for analysis of muscle function and histology. Verapamil-treated Ca_V_1.1^Δe29^/ClC-1^-/-^ mice showed significant weight gain that was equivalent to verapamil-treated WT mice (Fig 4B).

**Fig. 4.**
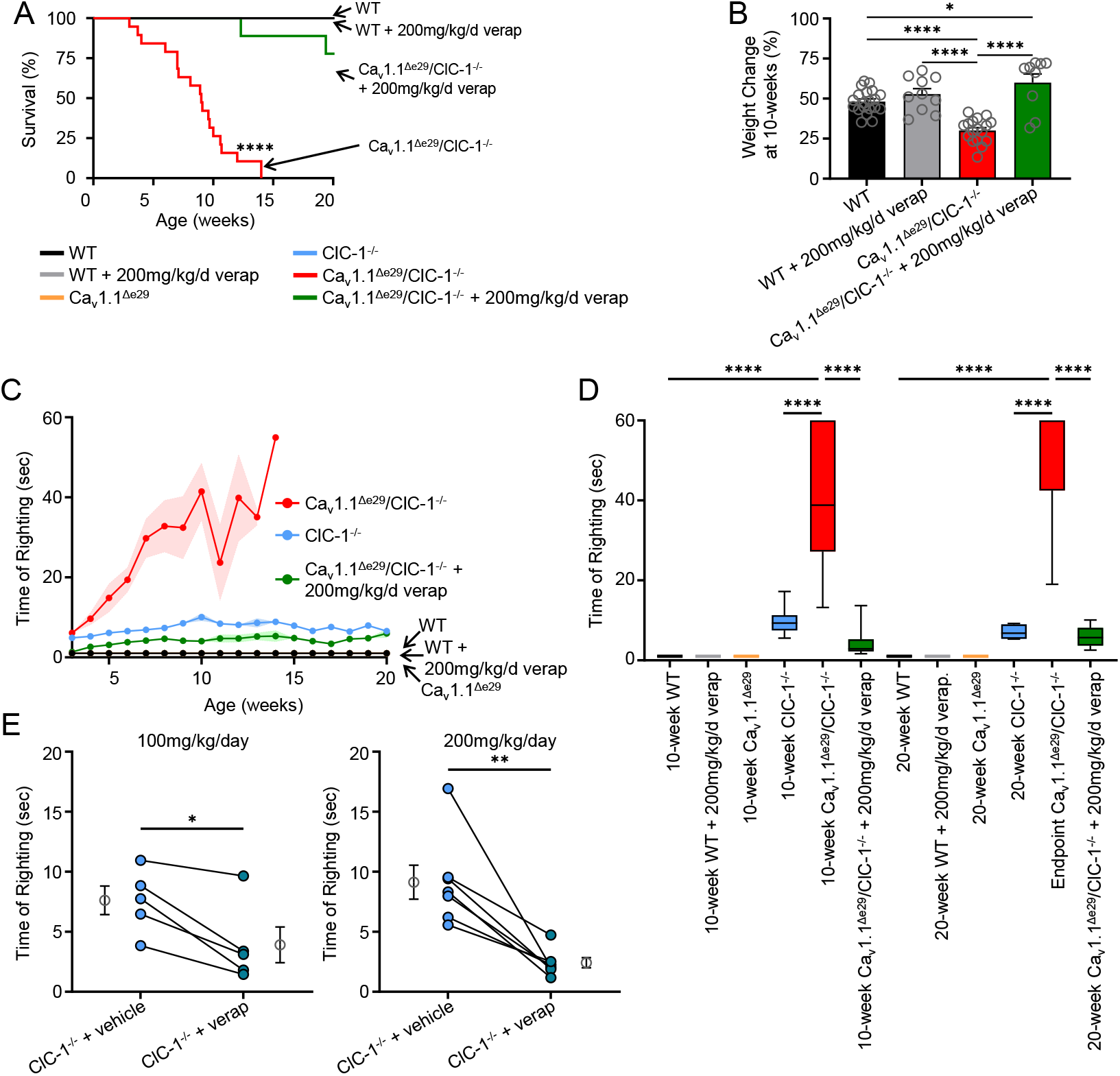
Verapamil treatment improves survival, body weight and motility of Ca_V_1.1^Δe29^/ClC-1^-/-^ mice. **A)** Kaplan-Meier survival analysis of WT (n=10; female=5, male=5), WT + 200mg/kg/day verapamil (n=10; female=5, male=5), Ca_v_1.1^Δe29^/ClC-1^-/-^ (n=19, female=9, male=10), and Ca_v_1.1^Δe29/Δe29^/ClC-1^-/-^ + verapamil (n=9, female=5, male=4). Verapamil is dosed in mouse nutrition/hydration food cups. Note: Log-Rank analysis of Ca_v_1.1^Δe29^/ClC-1^-/-^ vs. WT, WT + 200mg/kg/day verap, and Ca_v_1.1^Δe29^/ClC-1^-/-^ + 200mg/kg/day verap; P = <0.0001, <0.0001, and <0.0001 respectively. **B)** Percent body weight change from weaning at 10-wks of WT (n=20; female=10, male=10), WT + 200mg/kg/day verapamil (n=10; female=5, male=5), Ca_v_1.1^Δe29^/ClC-1^-/-^ (n=35, female=14, male=21), and Ca_v_1.1^Δe29/Δe29^/ClC-1^-/-^ + verapamil (n=9, female=5, male=4) **C)** Weekly time of righting reflex analysis of Ca_v_1.1^Δe29^/ClC-1^-/-^ (red), ClC-1^-/-^ (light blue), Ca_v_1.1^Δe29^ (orange), WT (black), Ca_v_1.1^Δe29^/ClC-1^-/-^ + verapamil (green) and WT + verapamil (grey) mice, **D)** Average time of righting reflex in vehicle and verapamil treated mice at 10- and 20-wks of age. Note: Untreated Ca_v_1.1^Δe29^/ClC-1^-/-^ mice do not survive to 20-wks of age, therefore the last recording before death is documented. Box notes Q1, median, and Q3; whiskers show minimum and maximum. **C, D)** WT (n=10; female=5, male=5), WT + 200mg/kg/day verapamil (n=10; female=5, male=5) Ca_v_1.1^Δe29^ (n=8; female=4, male=4), ClC-1^-/-^ (n=19; female=9, male=10), Ca_v_1.1^Δe29^/ClC-1^-/-^ (n=15; female=7, male=8), 10-wk Ca_v_1.1^Δe29/Δe29^/ClC-1^-/-^ + verapamil (n=16, female=7, male=9), and 20-wk Ca_v_1.1^Δe29/Δe29^/ClC-1^-/-^ + verapamil (n=7, female=4, male=3). **E)** Paired before (light circles) an after (dark circles) 2-wk verapamil treatment of ClC-1^-/-^ mice at 100mg/kg/day (left; n=5; female=3, male=2) and 200mg/kg/day (right; n=6; female=3, male=3) dosing in nutrition/hydration food cups. Symbols, closed circles, individual mice; open circles, means ± SEM. Statistical analysis of results in Fig. 3. are found in Supplementary Notes.

To assess the effect of verapamil on general mobility in Ca_V_1.1^Δe29^/ClC-1^-/-^ mice we again utilized the righting response. Chronic administration of verapamil significantly improved the righting response of Ca_V_1.1^Δe29^/ClC-1^-/-^ mice (Fig. 4C, green symbols) to near-WT levels (Fig. 4A, black symbols). Average righting times for 10 and 20 weeks of age (or last TRR trial before death) are presented in Fig. 4D. Surprisingly, Ca_V_1.1^Δe29^/ClC-1^-/-^ mice treated with verapamil showed a modest improvement of righting response over untreated ClC-1^-/-^ mice. Since acute exposure to verapamil had also reduced transient weakness and myotonia in single-channelopathy ClC-1^-/-^ muscle and 9-AC-treated WT muscle *ex vivo* (see above), in the context of WT (+e29) Ca_V_1.1, we also examined ClC-1^-/-^ mice before and after two-weeks of 100 and 200 mg/kg/d verapamil administration by oral feeding. The righting response improved at both doses (Fig. 4E), with 200 mg/kg/d treatment having the greater effect.

### Long-term verapamil administration improves respiratory and diaphragm function in Ca_V_1.1^Δe29^/ClC-1^-/-^ mice

Respiratory failure is a common complication of DM1 in its later stages(*29*). We therefore used whole-body plethysmography to assess respiratory function in bi-channelopathy mice. As compared to WT controls, the peak inspiratory flow rate (PIFR), peak expiratory flow rate (PEFR), and tidal volume (TV) were all reduced in Ca_V_1.1^Δe29^/ClC-1^-/-^ mice at 10 weeks of age (Fig. 5A,C,E). Long-term administration of verapamil rescued the deficits of PIFR, PEFR and TV to values similar to WT mice at ages 10 and 20 weeks (Fig. 5A-F). Since diaphragm muscle is preferentially affected in DM1, we investigated if respiratory impairment in Ca_V_1.1^Δe29^/ClC-1^-/-^ mice is accompanied by reduced diaphragm function. We examined the force-frequency relationship using strips of diaphragm muscle isolated from 10-week old Ca_V_1.1^Δe29^/ClC-1^-/-^ mice, and found a marked reduction of tetanic force compared to WT mice at frequencies >20 Hz (Fig. 5H). Remarkably, the reduction in diaphragmatic force was abrogated by chronic administration of verapamil (Fig. 5H), even though verapamil was not added to the bath during the force measurements. This correction of diaphragm force production was maintained at 20-weeks of age in Ca_V_1.1^Δe29^/ClC-1^-/-^ mice, with peak tetanic force production similar to WT mice (Fig. 5I). Representative tetanic force traces from 10- and 20-week old mice (Fig. 5G) demonstrated no overt difference of shortening velocity between the groups, however relaxation was markedly prolonged in diaphragm muscle isolated from verapamil treated Ca_V_1.1^Δe29^/ClC-1^-/-^ mice, indicating that anti-myotonia effects were ‘washed out’ during the force measurements, and supporting a myoprotective effect of chronic exposure to verapamil *in vivo*. Taken together, the rescue of diaphragm muscle function by verapamil supports the observed improvement of respiratory function. We next wanted to determine if verapamil had myoprotective effects on limb muscle. Indeed, we found that EDL muscles isolated from Ca_V_1.1^Δe29^/ClC-1^-/-^ mice that received long-term administration of 200mg/kg/d verapamil had significantly increased specific force production upon tetanic stimulations at 10-weeks (Supplemental Fig. S15A,B), and was sustained at 20-weeks of age (Supplemental Fig. S15C,D). Further, the muscle weight and cross-sectional area (CSA) of dissected 10-week old Ca_V_1.1^Δe29^/ClC-1^-/-^ EDL muscles were significantly smaller compared to those of age-matched WT mice (Supplemental Fig. S15E), suggesting that the muscles were atrophic. Verapamil treated Ca_V_1.1^Δe29^/ClC-1^-/-^ mice showed a significant increase in weight and CSA of isolated EDL muscles at 10 weeks that did not differ from age matched WT mice at 10 (Supplemental Fig. S15E) or 20 weeks (Supplemental Fig. S15F).

**Fig. 5.**
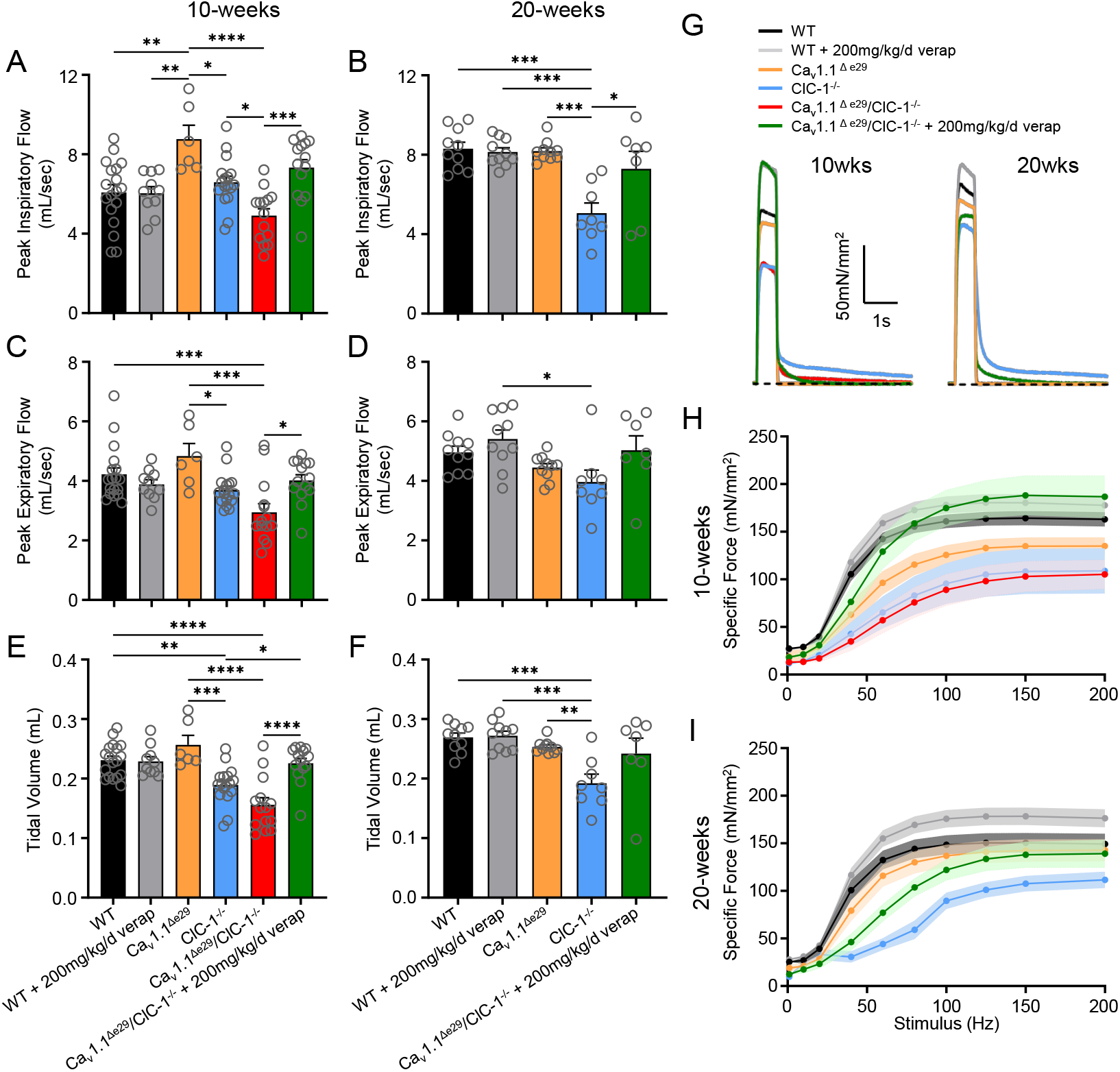
Verapamil treatment significantly improves respiratory function and diaphragm strength in Ca_v_1.1^Δe29^/ClC-1^-/-^ mice. Whole-body plethysmography for **A, C** and **E)** 10-wk (WT (n=18; female=9, male=9), WT + 200mg/kg/day verapamil (n=10; female=5, male=5) Ca_v_1.1^Δe29^ (n=6; female=3, male=3), ClC-1^-/-^ (n=17; female=8, male=9), Ca_v_1.1^Δe29^/ClC-1^-/-^ (n=14; female=7, male=7), and Ca_v_1.1^Δe29/Δe29^/ClC-1^-/-^ + verapamil (n=14, female=7, male=7)) and **B, D** and **F)** 20-wk old (WT (n=10; female=5, male=5), WT + 200mg/kg/day verapamil (n=10; female=5, male=5), Ca_v_1.1^Δe29^ (n=10; female=4, male=6), ClC-1^-/-^ (n=8; female=4, male=4), Ca_v_1.1^Δe29^/ClC-1^-/-^ + verapamil (n=7; female=4, male=3) mice in the indicated genotype and treatment groups. **A, B)** Peak inspiratory flow (mL/sec) and **C, D)** peak expiratory flow (mL/sec) of respiration. **E, F)** tidal volume (mL) of respiration. **G)** Representative tetanic (150Hz, 500ms) specific force traces from diaphragm strips isolated from 10-wk (left) and 20-wk (right) old mice in the indicated genotypes and treatment groups. **H** and **I)** Plot of average stimulation frequency dependence of specific force generated from diaphragm strips isolated from **H)** 10-wk (“n”=individual EDLs; WT (n=10; female=5, male=5), WT + 200mg/kg/day verapamil (n=6; female=3, male=3) Ca_v_1.1^Δe29^ (n=5; female=3, male=2), ClC-1^-/-^ (n=5; female=2, male=3), Ca_v_1.1^Δe29^/ClC-1^-/-^ (n=7; female=4, male=3), and Ca_v_1.1^Δe29/Δe29^/ClC-1^-/-^ + verapamil (n=7, female=3, male=4)) and **I)** 20-wk old (“n”=individual EDLs; WT (n=8; female=4, male=4), WT + 200mg/kg/day verapamil (n=8; female=4, male=4), Ca_v_1.1^Δe29^ (n=7; female=4, male=3), ClC-1^-/-^ (n=7; female=3, male=4), Ca_v_1.1^Δe29^/ClC-1^-/-^ + verapamil (n=7; female=4, male=3)) mice in the indicated genotypes and treatment groups. Symbols, open circles, individual mice; bars and closed circles, means ± SEM. Note: Untreated Ca_v_1.1^Δe29^/ClC-1^-/-^ mice do not survive to 20-wks of age. Statistical analysis of results in Fig. 5. are found in Supplementary Notes.

## Discussion

ClC-1 and Ca_V_1.1 channels are both required for normal function of skeletal muscle, and both undergo perinatal transitions in isoform expression. The predominant ClC-1 isoform expressed in fetal muscle includes a cryptic exon 7a, causing a shift of reading frame, nonsense-mediated decay of mRNA, truncation of channel protein, and loss of ion channel activity(*2, 11*). As fibers develop and the transverse-tubule system matures, the alternative splicing shifts to exclude exon 7a under the influence of MBNL, leading to high levels of chloride conductance. The chloride conductance then acts as an electrical buffer to stabilize the transmembrane potential during muscle activity(*30-32*). In recessive myotonia congenita (RMC), loss of ClC-1 is known to cause depolarization of the surface membrane during muscle activity, involuntary runs of action potentials (myotonia), and reversible loss of excitability (transient weakness).

Ca_V_1.1 is the voltage-sensor for ECC. Upon muscle excitation, Ca_V_1.1 undergoes a conformation change that triggers Ca^2+^ release from intracellular stores through physical coupling to RyR1. In fetal muscle exon 29 is skipped in around half the Ca_V_1.1 mRNAs, encoding Ca_V_1.1^Δe29^ protein that has substantial Ca^2+^ conductance(*12, 33*). The postnatal splicing of Ca_V_1.1 shifts to nearly 100% production of Ca_V_1.1^+e29^, again driven by MBNL, which has much lower Ca^2+^ conductance(*34*). Interestingly, the Ca^2+^ conductance of Ca_V_1.1^+e29^ is dispensable for ECC of mature muscle(*25*), but does contribute to fiber depolarization and transient weakness when ClC-1 is absent(*20*). Further, when the switch to Ca_V_1.1^+e29^ is entirely prevented by homozygous deletion of e29 in mice, the life-long expression of Ca_V_1.1^-e29^ leads to changes of fiber type specification and age-dependent signs of mitochondrial toxicity(*9*).

The CTG expansion in DM1 is highly unstable in somatic cells, causing age-dependent growth of the expanded repeat at different rates in different tissues. When size of the repeat and extent of toxic RNA reaches a threshold level in skeletal muscle, splicing of ClC-1 and Ca_V_1.1 reverts to fetal isoforms(*33*), bringing the circumstance of chloride channel myotonia together with increased Ca^2+^ entry through Ca_V_1.1^Δe29^. The main finding of our study is that combined Ca_V_1.1/ClC-1 bi-channelopathy is highly deleterious, causing aggravated myotonia and transient weakness, fixed weakness, respiratory impairment, and marked reduction of lifespan. It remains possible, if not likely, that other ECC-related splicing defects, such as expression of RyR1^-e70^ and Serca1a^-e22^, may also contribute, but our results suggest a central role for Ca_V_1.1/ClC-1 bi-channelopathy in the functional impairment of muscle.

Extrapolation of our findings to humans is not straightforward due to selective patterns of muscle involvement in DM1, effects of DM1 on expression of other genes, and limited survival of bi-channelopathy mice. It is important to note that other splicing defects, including *BIN1* and *DYS* have been previously implicated in DM1 myopathy(*8, 35*). It is therefore clear that Cl^-^/Ca^2+^ bi-channelopathy does not provide a unitary explanation for DM1 myopathy. To further dissect the contributory role of Cl^-^/Ca^2+^ bi-channelopathy, one approach in future studies is to subtract the influence of Ca_V_1.1^Δe29^ from the overall DM1 milieu using Ca^2+^ blockers, however this awaits the development of authentic models with longer repeats that fully replicate RNA toxicity observed in DM1 patients. Another important difference is that bi-channelopathy in our model is constitutive throughout all skeletal muscles, whereas effects of DM1 on different muscles are highly selective. For example, the myotonia in DM1 is often quite severe in distal limb and oromandibular muscles, which now can be attributed to combined ClC-1 loss and Ca_V_1.1 e29 skipping in those muscles, whereas in proximal limb muscles the myotonia and weakness is much less apparent. The mechanisms underlying selective muscle involvement are currently unknown, but evidence supports the idea that the extent splicing dysregulation is associated with degree of muscle weakness within and between patients (*10*). It is reasonable to expect that observations from bi-channelopathy mice may apply most directly to DM1 muscles that exhibit near-complete loss of ClC-1 and levels of Ca_V_1.1 e29 skipping approaching or exceeding heterozygous Ca_V_1.1^Δe29^ mice, and this clearly does occur in distal limb muscles that are preferentially affected, such as tibialis anterior (*10*). Finally, the duration of Cl^-^/Ca^2+^ channelopathy is limited by short survival of bi-channelopathy mice in this study. Therefore, in future studies it will be important to examine effects of Cl^-^/Ca^2+^ bi-channelopathy over longer intervals in specific muscle groups using conditional Ca_V_1.1^Δe29^ mice.

A limitation of our study is that cause of death in bi-channelopathy mice is not clearly defined. Almost certainly it results from skeletal muscle, since ClC-1 and Ca_V_1.1 are both expressed almost exclusively in this tissue. Since the animals do not show normal weight gain and exhibit diaphragmatic weakness, it is likely that inanition and respiratory insufficiency may both contribute.

Our results indicate a complex physiologic interplay of Cl^-^ and Ca^2+^ channels, in which increased Ca^2+^ entry through Ca_V_1.1^Δe29^ aggravates the myotonia and transient weakness from Cl^-^ channelopathy, and conversely that Cl^-^ channel myotonia may aggravate myopathy caused by excessive Ca^2+^ entry through Ca_V_1.1^Δe29^ channels. To a surprising extent, it appears that this chain of events can be broken by single intervention at the level of Ca^2+^ channels. Verapamil blockade of Ca_V_1.1 showed striking benefits for survival and respiratory function in bi-channelopathy mice, and also improved the myotonia and transient weakness in bi-channelopathy muscle fibers. In fact, the latter effects were even observed in single-channelopathy ClC-1^-/-^ fibers. Whether verapamil may also alleviate the long-term mitochondrial toxicity in single- or bi-channelopathy Ca_V_1.1^Δe29^ mice remains to be determined. Interestingly, Grant and colleagues reported in 1987 that another Ca^2+^ channel blocker, nifedipine, had anti-myotonia effects in DM1(*36*). To our knowledge, confirmatory studies were never carried out, possibly because genes for non-dystrophic myotonia were discovered soon thereafter and shown to encode sodium or Cl^-^ channels, undermining a potential connection to Ca^2+^ channels.

We caution against direct application of our results to the treatment of DM1 patients. The bi-channelopathy mice are not expected to exhibit any of the cardiac features of DM1, such as disease of the conduction system, which is potentially exacerbated by Ca^2+^ channel blockers. Further work is needed to identify agents that benefit Ca_V_1.1^Δe29^ channelopathy without aggravating conduction system disease or the tendency for low blood pressure. This includes the possibility, which to our knowledge has not been a priority previously, of developing agents that are selective for Ca_V_1.1 channels in skeletal muscle over Ca_V_1.2 channels in cardiac and smooth muscle.

## Supporting information

Statistical Analysis for Results in Main Text Figures

Statistical Analysis for Results in Supplemental Figures

## Study approval

All procedures in this study complied with the University of Rochester Committee on Animal Resources.

## Acknowledgements

We thank the members of the Lueck Laboratory and Dr. Amy Martin for reading and editing the manuscript and constructive discussion throughout the study. We thank Victoria Shwe, Julia Hyatt, Linda Richardson and Sara Leistman for their technical assistance. We thank Dr. Terry W. Wright for help with initial plethysmography measurements and Kevin J. Potcner, Statistical Scientist from JMP Statistical Discovery^®^, LLC for guidance with statistical analyses. We gratefully acknowledge the Genome Editing Core at Augusta University for generating the SERCA1^Δe22^ and Ca_V_1.1^Δe29^ mice mouse models necessary to support this research. We thank Dr. Kevin P. Campbell for providing RyR1^Δe70^ mice, which were generated at the University of Iowa Genome Editing Core Facility directed by William Paradee, PhD and supported in part by grants from the NIH and from the Roy J. and Lucille A. Carver College of Medicine. We wish to thank Norma Sinclair, Rongbin Guan and Joanne Schwarting for their technical expertise in generating transgenic mice. We thank Dr. Jacqueline Heller and The Klein Family for their generous support.

## Funding

National Institute of Health Grant T90DE021985 (LAC)

National Institute of Health Grant T32 GM135134 (LAC)

Doctoral Fellowship from The Myotonic Dystrophy Foundation (LAC)

National Center for Advancing Translational Sciences of the National Institutes of Health UL1 TR002001 (JDL)

National Institute of Health Grant R01 AR079424 (JDL)

## Author contributions

Conceptualization: LAC, MTS, CAT, JDL Methodology: JDL

Investigation: LAC, MTS, KME, JDL Funding acquisition: LAC, JDL

Project administration: JDL

Supervision: JDL

Writing – original draft: LAC, MTS, CAT, JDL

Writing – review & editing: LAC, MTS, KME, CAT, JDL

## Competing interests

J.D.L., L.A.C. and C.A.T. are co-inventors of the technology presented in this study. PCT/US22/79593 Claims Priority to U.S. Provisional Patent Application Nos. 63/277,816 & 63/374,378, Title: TREATMENT OF MYOTONIC DISORDERS. Inventors - University of Rochester School of Medicine and Dentistry – Inventors - J.D.L., L.A.C. and C.A.T. pertains to the use of verapamil to treat myotonic disorders in this study.

## Data and materials availability

All data are available in the main text or the supplementary materials.

## Notes

### Competing Interest Statement

The authors have declared no competing interest.

