## Supplemental Methods and Figures for "Combinatorial chloride and calcium channelopathy in myotonic dystrophy"

**The PDF file includes:**

Materials and Methods
Figs. S1 to S15
Tables S1 to S4

**Other Supplementary Materials for this manuscript include the following:**

Statistical Analysis Figs. 1 to 5
Statistical Analysis Figs. S7 to S9 and S11, S13 and S14

#### Materials and Methods

##### Generation of mouse lines

CRISPR/Cas9 was used to generate forced exon deletion mice. Exon 29 of CACNA1s, exon 22 of SERCA1, and exon 70 of RyR1 were removed by targeting ~50 nucleotides into intronic sequence on either side of the intron/exon boarder, to ensure that both the exon and the splice donor/acceptor sequence were removed. Founder lines were crossed to C57bl6 (Jackson Laboratories) >6 generations to eliminate possible off-target mutations. sgRNA sequences used to generate mice are listed in Table S1. The removal of exon 29 in $Ca_v1.1$, exon 22 in SERCA1, and exon 70 in RyR1 was validated by RNA isolation of tibialis anterior and Sanger sequencing of RT-PCR products between exons noted for each transcript in Table S2. Primers used for Sanger sequencing are also noted in Table S2. To generate $Ca_v1.1^{\Delta e29}$/ClC-$1^{-/-}$, SERCA$1^{\Delta e22}$/ClC-$1^{-/-}$, and RyR$1^{\Delta e70}$/ClC-$1^{-/-}$ mice, *adr-mto2J* mice (Jackson Laboratory), which have a frameshift mutation in *Clcn1* (ClC-$1^{-/-}$) causing a recessive generalized chloride-channel myotonia, were bred with $Ca_v1.1^{\Delta e29/\Delta e29}$, SERCA$1^{\Delta e22/\Delta e22}$, and RyR$1^{\Delta e70/\Delta e70}$mice(*11*). A*dr-mto2J* were obtained from Jackson Laboratory and bred to achieve congenic C57bl6/J strains. WT C57bl6/J controls were obtained from Jackson Laboratory when littermates could not be used. All mice were housed, bred, cared for, and experimented on in accordance with University of Rochester Committee on Animal Resources approval. For $Ca_v1.1^{\Delta e29}$/ClC-$1^{-/-}$ mice, terminal endpoints were determined by age of the mice or when time of righting reflex was >60 seconds.

##### Verapamil administration

Mice were provided verapamil via ingestion of Nutra-Gel Complete Nutrition food (Bio-Serv) to a final dose of ~200mg/kg/day verapamil (V4629, (±)-Verapamil hydrochloride, Sigma-Aldrich). Food cups contained 0.1% W/W verapamil, which was first solubilized in 1mL of $dH_2O$. Mice that did not receive verapamil, were provided Nutra-Gel Complete Nutrition food that had 1ml of $dH_2O$ added to serve as vehicle control. Mice were provided food cups immediately after weaning and food was changed daily and weighed for consumption. No other food source was provided. Water bottles were provided and changed weekly.

##### Body weight measurements

Body weight of the mice was measured every other day following weaning. For statistical analysis, two-way ANOVA was performed on the average of each week's measurements. Percent body weight change at 10-weeks was determined with the first weight measurement taken post weaning and the body weight reached at 10-weeks of age. For this data, a one-way ANOVA with multiple comparisons was performed.

##### Time of righting reflex measurements

Time of righting reflex was tracked weekly after the start of verapamil treatment. Mice were held in a supine position on a level surface then released. Two persons blinded to genotype and treatment began timing immediately after release, and timing was stopped once all four paws were down on the surface. This was repeated for a total of 10 timed righting responses with a minimum of five min in between each recording to avoid the warm-up phenomenon(*15*). Recordings less than one second were recorded as "1" and righting was considered a failure if righting took more than 60 sec. If failure was reached, no further recordings were measured for that mouse for that week. The two longest and two shortest recording for each mouse was removed and the average of the middle 6 recording were used for analysis. We performed two-way ANOVA on the week-to-week results, and one-way ANOVA with multiple comparisons for the timepoint data (10-week and 20-week).

##### Whole body plethysmography

Whole body plethysmography (Buxco® Small Animal Whole Body Plethysmography, Data Science International) was used to measure respiratory function at 10-weeks (all groups) and 20-week (excluding untreated $Ca_v1.1^{\Delta e29}$/ClC-$1^{-/-}$). Mice were acclimated to the system for two days with data being collected on day three. For all three days, single mice were placed conscious and unrestrained in WBP chamber and subjected to a 15 min acclimation period, followed by 10 min of data acquisition(*37*). Average data from the 10 min acquisition period were analyzed by one-way ANOVA with multiple comparisons.

**Whole-cell patch-clamp recording of calcium currents from dissociated adult FDB fibers**

Mice at approximately 4-weeks postnatal were sacrificed by cervical dislocation and decapitation. Isolated *flexor digitorum brevis* (FDB) muscle fibers for electrophysiology experiments were obtained by previously described methods(*4*). Briefly, FDB muscles were immediately micro-dissected submerged in standard electrophysiology Ringer's solution (146mM NaCl, 5mM KCl, 2mM $CaCl_2$, 1mM $MgCl_2$, and 10mM HEPES at pH 7.4 w/ NaOH). Excised muscles were then digested for 1-hour in an oscillating water bath at 37°C in Ringer's solution supplemented with 2 mg/mL Collagenase A (Roche, Mannheim, Germany). Following enzymatic digestion, the muscles were mechanically dissociated by trituration with fire-polished Pasteur pipettes of decreasing bore diameter to obtain isolated single FDB fibers. Fibers were then plated on 35mm plastic cell-culture dishes for experiments. All patch-clamp experiments were completed within 8-hours of the sacrifice of the mouse. To record $Ca_V1.1$ calcium currents with whole-cell patch-clamp, solutions were designed to isolate calcium currents (TTX (Cayman Chemical Company, Ann Arbor, MI) to block sodium-currents, TEA, 4-AP, and cesium to block potassium currents, and anthracene-9-carboxylic acid (9-AC) to block chloride currents). External recording solution was 145mM TEA-methanesulfonic acid, 10mM $CaCl_2$, 10mM HEPES, 2mM $MgSO_4$, 1mM 4-aminopyridine, 0.1mM 9-AC, and 0.001mM tetrodotoxin, pH 7.4 with TEA-OH. Low-resistance pipettes were fashioned from thin-walled borosilicate glass with a Sutter P-97 puller (Sutter Instruments, Novato, CA) and fire-polished to a resistance <1 MΩ. Pipettes were filled with an internal solution with 140mM Cs-aspartate, 10mM $Cs_2$-EGTA, 5mM $MgCl_2$, and 10mM HEPES, pH 7.4 with CsOH. Currents were recorded with standard voltage-step protocols with 500ms voltage steps from -50 to -80 to 80mV at 10mV intervals, as described in(*38*). Data was collected using an Axopatch 200B amplifier (Axon Instruments, San Jose, CA) and digitized with a Digidata 1550b (Axon Instruments). A -P/4 online subtraction protocol was utilized to correct linear components of leak and capacitive currents. To limit voltage error, whole-cell parameters were adjusted, and series resistance was compensated (90%). Recordings were sampled at 10kHz and filtered at 2kHz. Cell capacitance and other passive properties of the muscle fibers were either obtained via Clampex 10 (Molecular Devices) or calculated based on integration of a 10-mV voltage step from resting. Currents were normalized to fiber capacitance to obtain current densities (pA/pF). All experiments were completed at room temperature. Peak current densities obtained for the family of voltage steps were plotted and fit with the following equation to obtain I-V curves.

$$I=G\text{max}*(V-V\text{rev})/(1+\exp(-(V-V1/2)/kG))$$

In this equation, $I$ correspond to the peak current of a specific test potential normalized to cell capacitance, $G$max is the maximum calcium conductance, $V$1/2 is the half-maximal activation potential, $V$rev is the reversal potential, and $kG$ is the slope factor.

**Histology and fiber typing**

For hematoxylin and eosin (H&E) staining, as well as fiber typing by immunohistochemistry, tibialis anterior and diaphragm were isolated from 10-week and 20-week mice (n=5/group) and snap frozen. Transverse 10-micron cryosections from each muscle were collected on glass slides. Slides were submerged in hematoxylin for 3.5 min, followed by 1 wash in $ddH_2O$, 1% HCl, $ddH_2O$, ammonia water, $ddH_2O$, and submerged in eosin for 5 min. Slides were dehydrated by two 30 sec incubations in 95% ethanol, two 30 sec incubations in 100% ethanol, one 30 sec incubation followed by a 5 min incubation in Hemo-De (xylene substitute). The slides were then mounted with Permount (Fisher) and imaged on an Olympus BX53. Immunohistochemistry used to determine fiber type was performed as in as in Bachman J. et al. (*39*). Briefly, slides were permeabilized once with PBST (PBS with 0.2% Triton X-100) for 10 min then blocked for 30 min in 10% normal goat serum (NGS, Gibco) in 1X PBS at room temperature. After, slides were blocked for 60 min in 3% affinipure Fab fragment anti-mouse IgG (H&L) (Jackson Immuno-research)

(0.1mg/ml) with 2% NGS in 1X PBS at room temperature then washed three times for 10 min each in PBS. Slides were incubated for 14-18hrs at 4°C in primary antibodies (noted in Table S3). Next, slides were washed three times for 10 min 1X PBS at room temperature then secondary antibodies (see Table S3) were added and incubated for 60 min. After another set of three washes for 10 min in 1X PBS at room temperature, slides were mounted with Fluromount-G (Invitrogen). Slides were imaged using a Keyence BZ-X810 All-in-One Florescence Microscope (Leica).

***Ex vivo* muscle contraction**

Muscle strength and frequency dependence were measured by *ex vivo* muscle contraction performed on an Aurora Scientific 1200A *ex vivo* system equipped with 809B muscle testing system with a 300C-LR force transducer and 701C stimulator (Aurora Scientific). For this experiment, 10- and 20-week WT, WT + verapamil, $Ca_v1.1^{\Delta e29}$, $ClC\text{-}1^{-/-}$, untreated $Ca_v1.1^{\Delta e29}/ClC\text{-}1^{-/-}$ (only at 10-weeks), and $Ca_v1.1^{\Delta e29}/ClC\text{-}1^{-/-}$ + verapamil mice were used. Mice were anaesthetized by 2% inhaled isoflurane. For the EDL muscle, the proximal and distal tendons of the EDL were exposed after removal of the tibial anterior and tied using suture thread. The proximal tendon was set on an immobile post and the distal tendon hooked to the force transducer. For diaphragm, a 4mm strip was dissected from the right costal hemidiaphragm as in Hakim, C. et al. (2019)(*40*). Suture was used to connect the diaphragm strip to the stationary post from the ribs and the central tendon was secured to the force transducer. For both muscle groups, the muscles were submerged between platinum electrodes in warmed (30°C) and oxygenated (95% $O_2$ and 5% $CO_2$) Ringer's buffer (1.2mM $NaH_2PO_4$, 1mM $MgSO_4$, 4.83mM KCl, 137mM NaCl, 2mM $CaCl_2$, 10mM glucose, 24mM $NaHCO_3$ at pH 7.4)(*41*). EDL and diaphragm muscles were equilibrated for 10 min before determining optimal length ($L_o$) and supramaximal output (120% stimulating voltage)(*42*). Muscles were then subjected to a twitch warm-up protocol (three 200ms 1Hz stimuli separated by 20 sec), and tetanus warm up protocol (three 500ms, 150Hz stimuli separated by 1 min) before frequency dependence was determined. Frequency dependence was determined with increasing 500ms stimulations separated by one min. For EDL the stimulations were as follows, 1Hz (200ms), 25Hz (500ms onward), 50Hz, 75Hz, 100Hz, 125Hz, 150Hz, 175Hz, 200Hz, 250Hz. For diaphragm, stimulations were 1Hz (200ms), 10Hz (500ms onward), 20Hz, 40Hz, 60Hz, 80Hz, 100Hz, 125Hz, 150Hz, 200Hz. Muscle force was recorded using 610A Dynamic Muscle Control LabBook software (Aurora Scientific) and analyzed using Clampfit 10.7.0.3 software (Molecular Devices). Specific force was calculated using wet weight of the muscle and optimized length between the proximal and distal myotendinous junctions(*41, 43*). Two-way ANOVA analysis was used to determine statistical differences between groups for frequency dependence.

**Quantification of myotonia**

Myotonia was measured using *ex vivo* muscle contraction. For experiments on 20-week old mice, EDLs were isolated from WT and $Ca_v1.1^{\Delta e29}$ mice and optimized as in the previous section. EDLs were first equilibrated for 25 min in Ringer's solution. Muscles were then subjected to a protocol to determine myotonia with 3 successive 200ms 1Hz twitches separated by 20 sec, a 3 min rest period, then 3 successive 500ms 150Hz tetani separated by 3 min. Ringer's solution with 100μM 9-AC was then flowed into the bath for 25 min and the myotonia protocol was repeated. This timing ensured there as enough time for solution exchange and for the muscle to equilibrate in the new bath. To determine the impact of verapamil on myotonia in WT and $Ca_v1.1^{\Delta e29}$ muscle, the myotonia protocol (noted earlier in this section) was performed in the presence of Ringer's solution containing either 5μM or 20μM verapamil then followed by Ringer's solution containing 100μM 9-AC and 5μM or 20μM verapamil. For the 6-weeks-of-age experiments, EDLs were isolated from 6-week WT, $Ca_v1.1^{\Delta e29}$, $ClC\text{-}1^{-/-}$, $Ca_v1.1^{\Delta e29}/ClC\text{-}1^{-/-}$ mice and optimized as in the previous section. For pre-treatment recordings, muscles were equilibrated for 25 min in Ringer's solution and the myotonia protocol was performed. Ringer's solution containing 20μM verapamil was added to the bath using gravity flow and equilibrated for 25 min. The myotonia protocol was then repeated. Muscle force was recorded using 610A Dynamic Muscle Control LabBook software (Aurora Scientific) and traces were analyzed using Clampfit 10.7.0.3 software (Molecular Devices) to calculate area under the curve. The area under the curve was then normalized to the specific force (calculated as in the previous section) and used for statistical analysis.

**Transient weakness measurements**

Transient weakness was measured in EDL muscle with *ex vivo* muscle contraction. EDLs were isolated and optimized from 20-week-old WT or $Ca_v1.1^{\Delta e29}$ mice as mentioned in previous sections. The experiments were contralaterally controlled with one side equilibrated in normal Ringer's solution, and the contralateral muscle was equilibrated in Ringer's solution with 100µM 9-AC for at least 25 min. Muscles were then subjected to the transient weakness protocol (45 successive 500ms 100Hz tetani separated by 4 sec). To determine the impact of verapamil on transient weakness, verapamil was dissolved in Ringer's solution and delivered to the bath by gravity flow. Using a dual-bath system, one muscle group received verapamil and the contralateral muscle received verapamil + 9-AC. To determine verapamil's impact, we tested concentrations of both 5µM and 20µM and equilibrated for 25 min before performing the transient weakness protocol. For 6-weeks-of-age experiments, EDLs were isolated from 6-week WT, $Ca_v1.1^{\Delta e29}$, $ClC\text{-}1^{-/-}$, $Ca_v1.1^{\Delta e29}/ClC\text{-}1^{-/-}$ mice and equilibrated for 25 min in Ringer's media containing no verapamil or 20µM verapamil (contralaterally). The transient weakness protocol was then performed. Muscle force was recorded and analyzed as previously described. Specific force was calculated, and for statistical analysis, we also quantified percent of the initial force generated.

**Statistical Analysis**

All average data is represented as means ± S.E.M. Data points collected for a single mouse were represented as open circles (one mouse is n=1), however, for *ex vivo* muscle contraction techniques, open circles represent one muscle (one muscle is n=1). The number of sampled units, n, upon which we reported statistics, is the single mouse for the *in vivo* experiments (one mouse is n = 1). GraphPad Prism 9 software was used for statistical analyses. One-way or Two-way ANOVA with multiple comparisons were used to test for significant differences between each of the groups tested. For survival, each group was compared to each other group Kaplan-Meier log-rank test. For all figures, *P < 0.05, **P < 0.01, ***P < 0.001, and ****P < 0.0001 were used.

#### Fig. S1

A

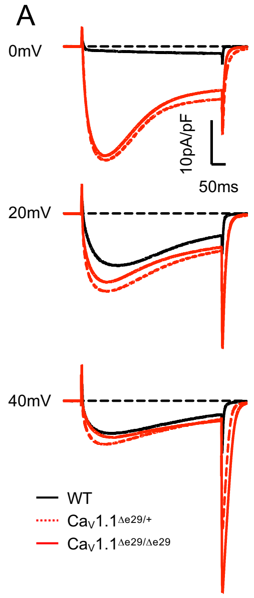


B

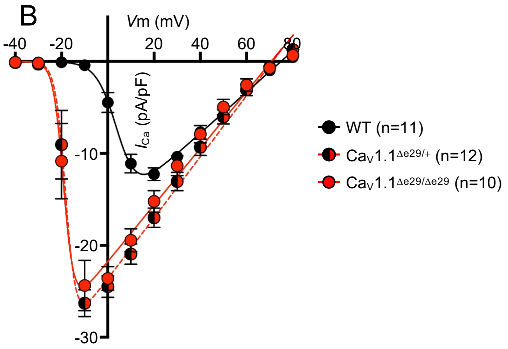


C

| Genotype: | $G_{max}$ (pS/pF) | $V_{rev}$ (mV) | $V_{1/2\ max}$ (mV) | $K_{act}$ (mV) |
|---|---|---|---|---|
| WT (C57Bl/6) | 0.224 ±0.011 | 74.897 ±1.485 | 4.568 ±0.758 | 4.350 ±0.530 |
| $Ca_V1.1^{\Delta E29/+}$ | 0.332 ±0.017 **** | 71.228 ±2.052 | -18.128 ±1.100 **** | 2.221 ±1.936 |
| $Ca_V1.1^{\Delta E29/\Delta E29}$ | 0.308 ±0.020 ** | 70.679 ±2.626 | -19.019 ±1.077 **** | 2.220 ±1.145 |
| Statistics (P-value) | <0.0001 | 0.2863 | <0.0001 | 0.3793 |

Table A: Electrophysiology – Modified Boltzmann Fitting Parameters
*All data presented mean ± SE; Standard One-Way ANOVA analysis P-value reported; * (stars) indicate significant differences compared to wildtype (2-sample t-test with Turkey's correction for multiple comparisons). No significant differences between homozygote ($Ca_V1.1^{\Delta e29/\Delta e29}$) and heterozygote ($Ca_V1.1^{\Delta e29/+}$) recordings were observed.*

D

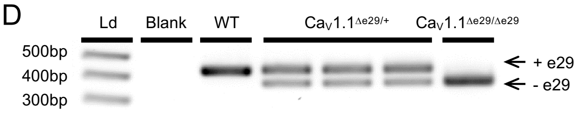


**Fig. S1. Heterozygous and homozygous $Ca_v1.1^{\Delta e29}$ mice exhibit similar $Ca_v1.1$ voltage-dependence and peak current densities in *flexor digitorum brevis* muscle. A)** Representative current density traces from whole cell patch clamp of *flexor digitorum brevis* fibers isolated from 4-week WT (black), $Ca_v1.1^{\Delta e29/+}$ (red, dashed), and $Ca_v1.1^{\Delta e29/\Delta e29}$ (red, solid) mice at 0 mV (top), +20mV (middle) and +40mV (bottom). **B)** Plot of average current-voltage relationship of $Ca_v1.1$ activity measured in WT (black), $Ca_v1.1^{\Delta e29/+}$ (red and black circles, red dashed), and $Ca_v1.1^{\Delta e29/\ \Delta e29}$ (red circle, solid red line) *flexor digitorum brevis* fibers isolated from 4-wk mice. **C)** Table of modified Boltzmann fitting parameters. **D)** RT-PCR products of $Ca_v1.1$ RNA isolated from *tibialis anterior* from 10-wk mice. PCR amplifications are from exons 27 to 31 of Cacna1s cDNA.

#### Fig. S2

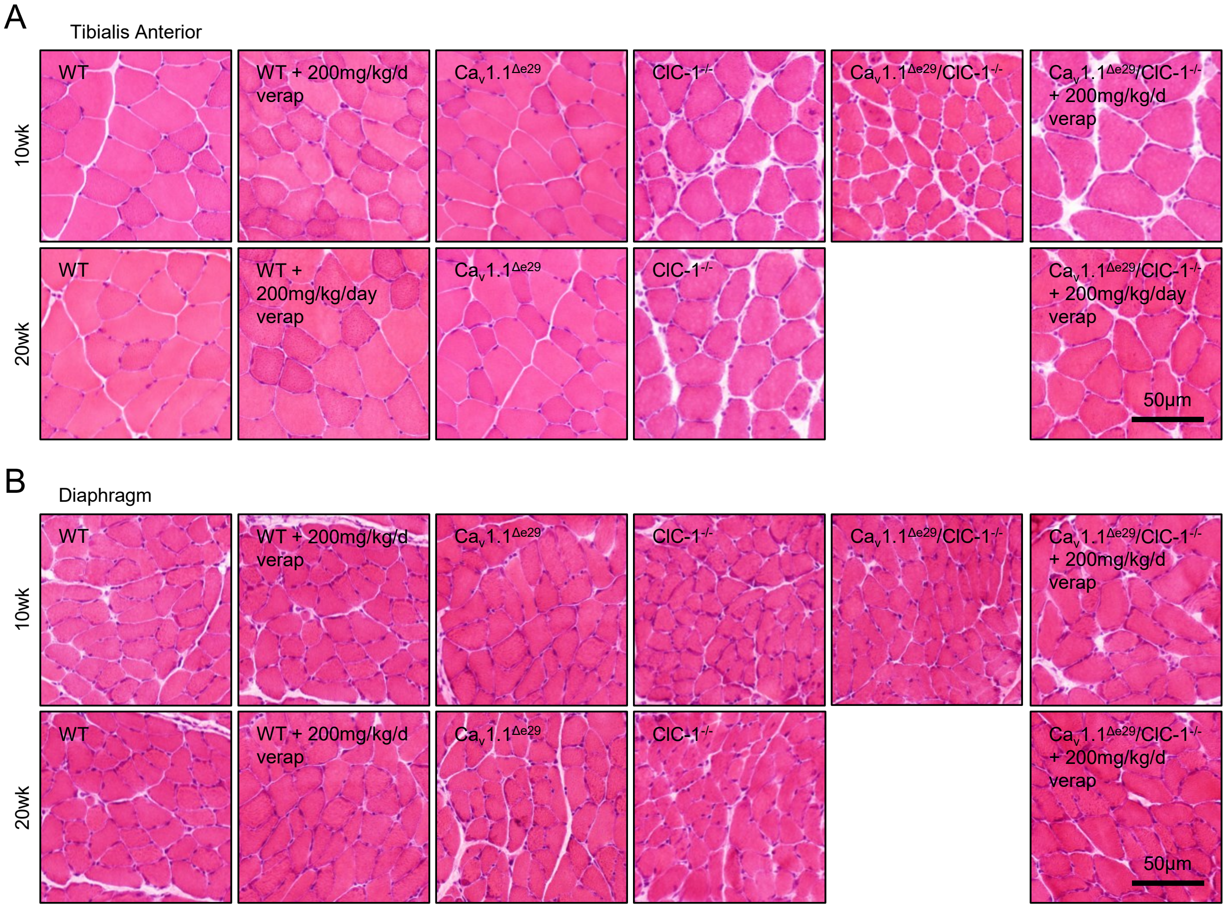


**Fig. S2. Ca$_v$1.1$^{\Delta e29}$/ClC-1$^{-/-}$ genotype does not impart overt dystrophic histological features in limb or diaphragm muscle. A)** Hematoxylin and Eosin staining of 10µm transverse sections of snap frozen tibialis anterior from 10-wk (top) and 20-wk (bottom) tissue samples. **B)** Hematoxylin and Eosin staining of 10µm transverse sections of snap frozen diaphragm from 10-week (top) and 20-week (bottom) tissue samples.

**Fig. S3**

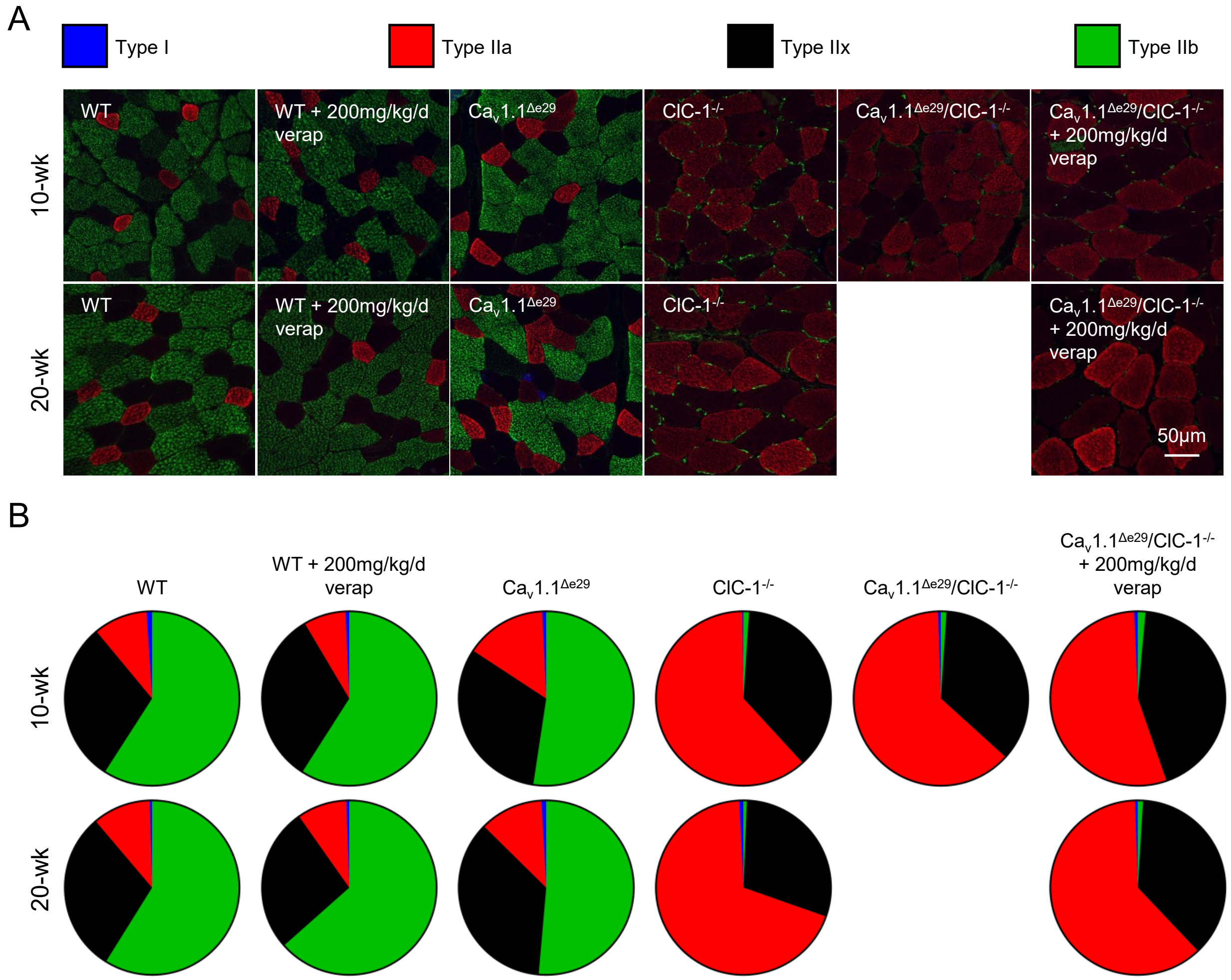


**Fig. S3. $Ca_v1.1^{\Delta e29}/ClC\text{-}1^{-/-}$ limb muscle fiber type distribution is not altered from $ClC\text{-}1^{-/-}$. A)** Representative images of fiber-type immunostaining for 10-wk (top) and 20-wk (bottom) tibialis anterior muscle isolated from indicated genotype and treatment groups. MYHC type IIB fibers, green; IIX, black; IIA, red; I, blue. Scale bars: 50µm. (n=5/group). **B)** Average quantification of fiber-type percentages of tibialis anterior muscle isolated from indicated genotype and treatment groups. Type IIB fibers (green), IIX, (black) IIA, (red) I, (blue). Scale bars: 50µm. (n=5/group)

**Fig. S4**

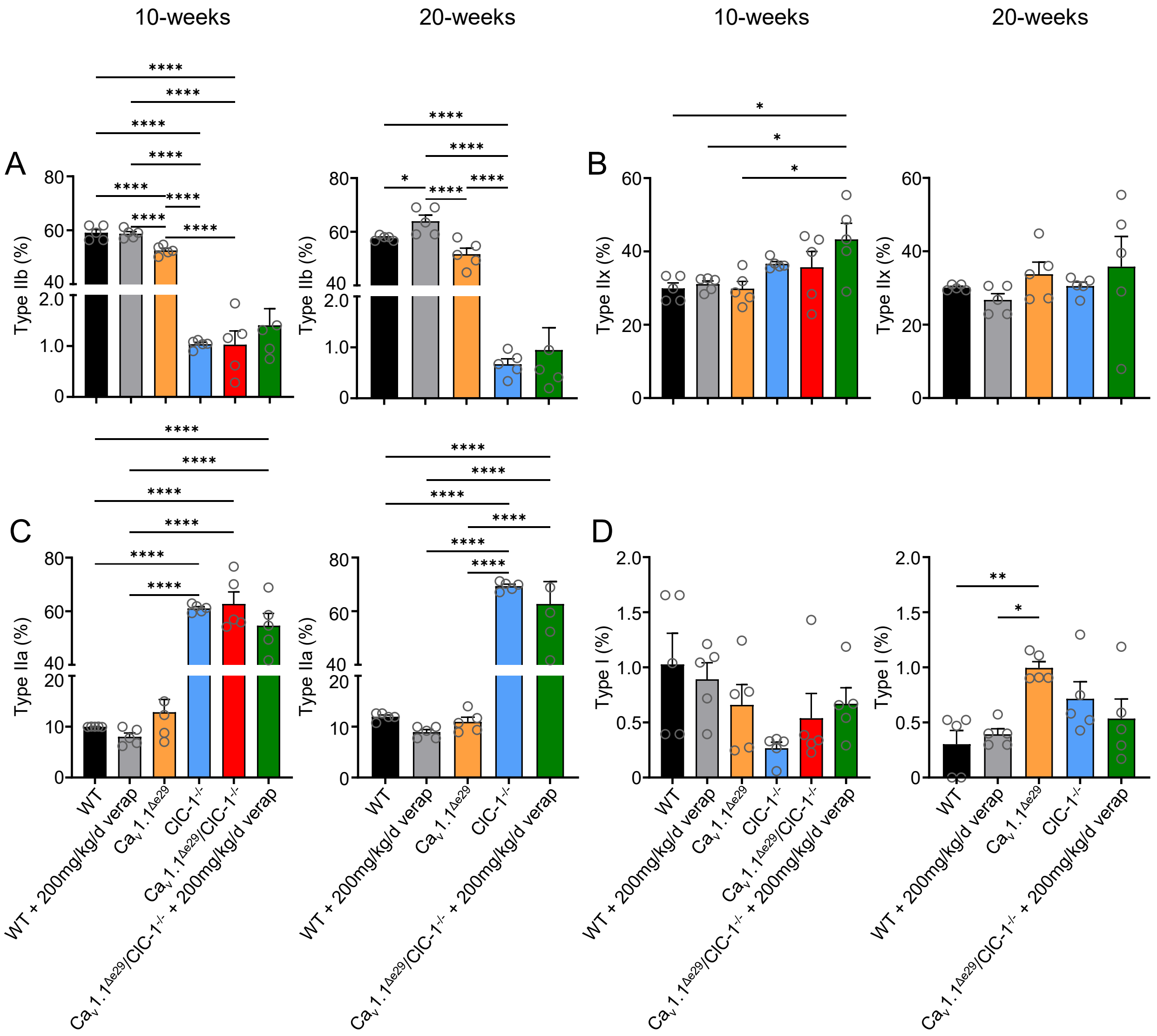


**Fig. S4. Quantification and statistical analysis of fiber type distribution of tibialis anterior muscle. A-D)** Quantification of **A)** type IIb **B)** type IIx **C)** type IIa **D)** type I fibers for 10-wk (left) and 20-wk (right) tibialis anterior (n=5/group). One-way ANOVA with multiple comparisons analysis performed, * = P < 0.05, ** = P < 0.01, *** = P < 0.001, and **** = P < 0.0001.

### Fig. S5

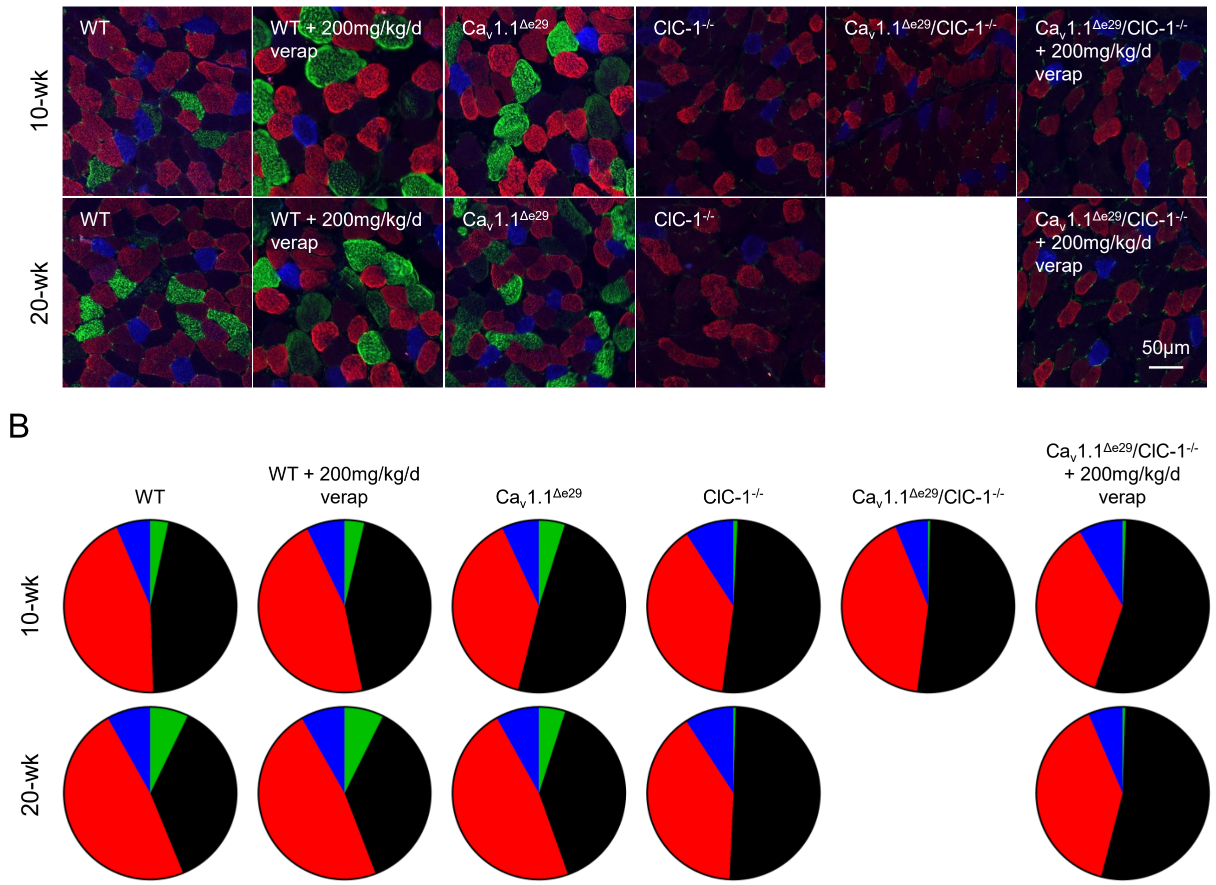


**Fig. S5. Ca$_v$1.1$^{\Delta e29}$/ClC-1$^{-/-}$ diaphragm muscle fiber type distribution is not altered from ClC-1$^{-/-}$. A)** Representative images of fiber-type immunostaining for 10-wk (top) and 20-wk (bottom) diaphragm muscle isolated from indicated genotype and treatment groups. MYHC type IIB fibers, green; IIX, black; IIA, red; I, blue. Scale bars: 50 µm. (n=5/group). **B)** Average quantification of fiber-type percentages of diaphragm muscle isolated from indicated genotype and treatment groups. Type IIB fibers (green), IIX, (black) IIA, (red) I, (blue). Scale bars: 50 µm. (n=5/group)

**Fig. S6**

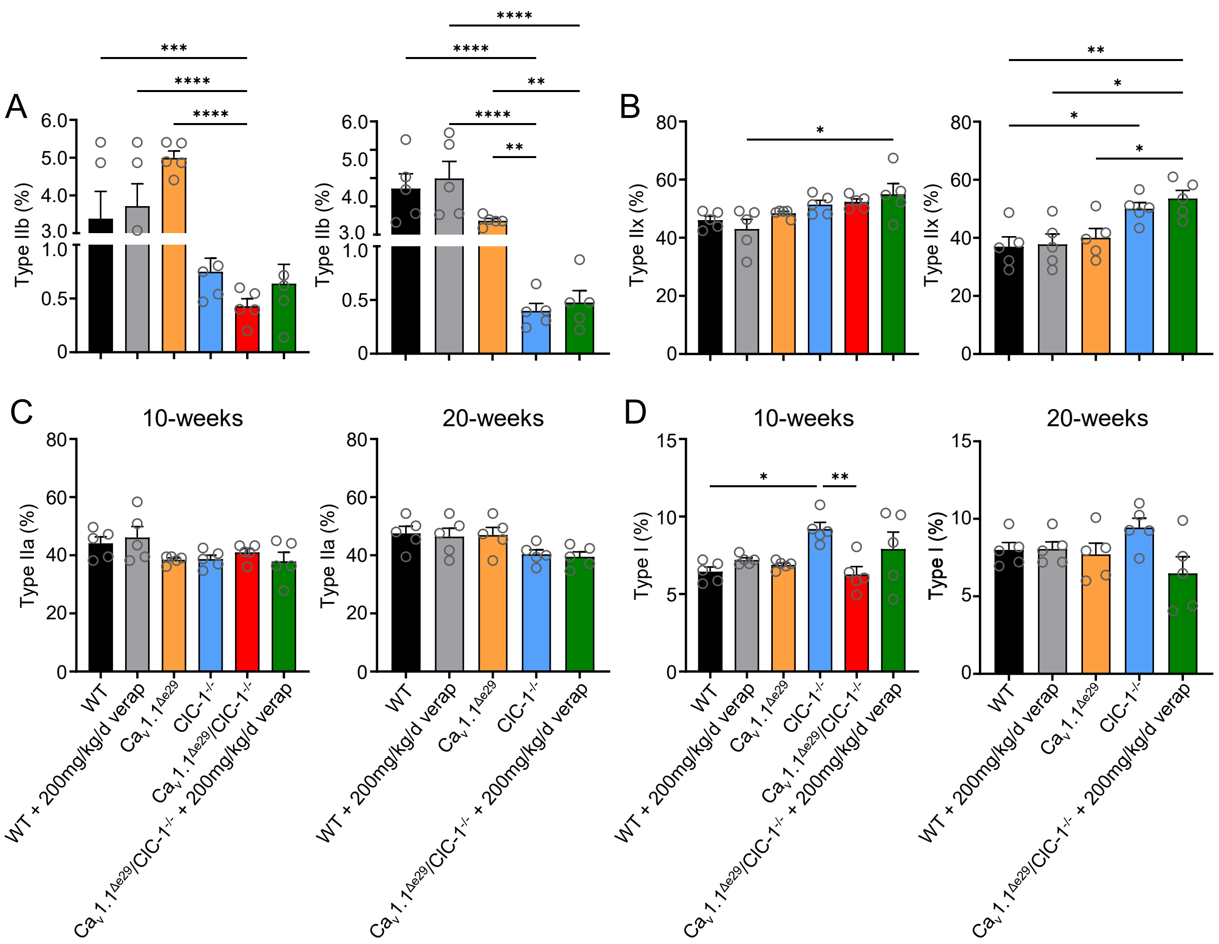


**Fig. S6. Quantification and statistical analysis of fiber type distribution of diaphragm muscle. A-D)** Quantification of **A)** type IIb **B)** type IIx **C)** type IIa **D)** type I fibers for 10-wk (left) and 20-wk (right) diaphragm (n=5/group). One-way ANOVA with multiple comparisons analysis performed, * = $P < 0.05$, ** = $P < 0.01$, *** = $P < 0.001$, and **** = $P < 0.0001$.

**Fig. S7**

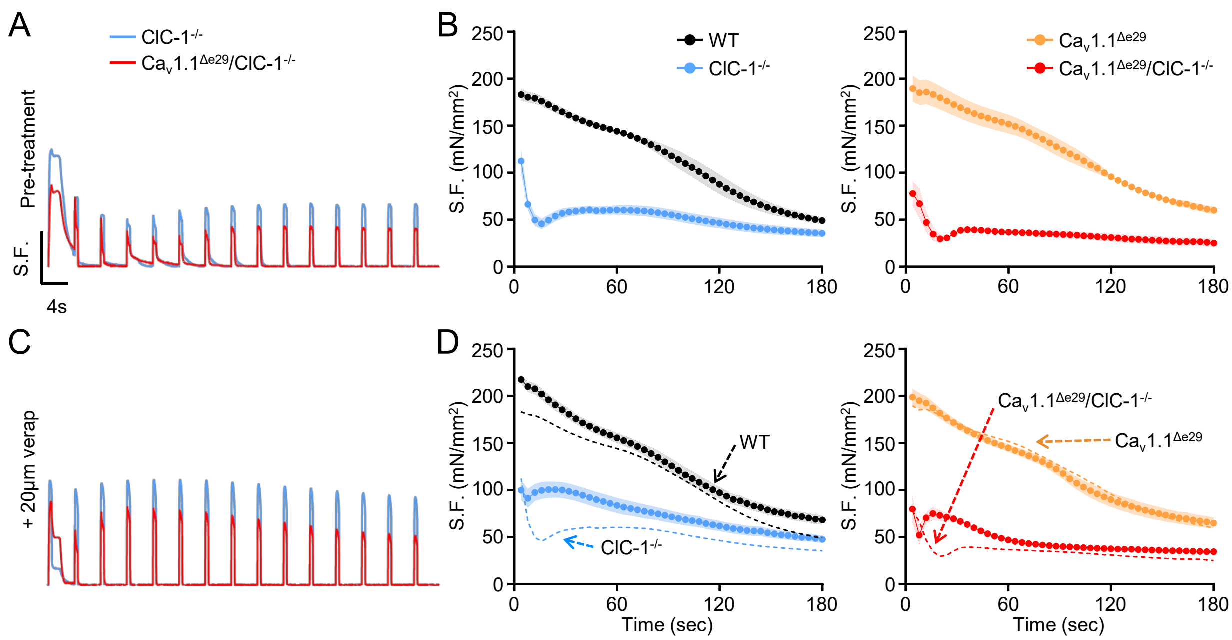


**Fig. S7. Ca$_v$1.1$^{\Delta e29}$/ClC-1$^{-/-}$ muscle exhibits severe transient weakness that is significantly improved by the addition of verapamil (not normalized to the first peak force). A, C)** Representative specific force traces of the first 15 tetani (100Hz, 500ms) separated by four seconds, recorded *ex vivo* from EDLs isolated from 6-wk ClC-1$^{-/-}$ (blue) and Ca$_v$1.1$^{\Delta e29}$/ClC-1$^{-/-}$ (red) mice in the **A)** absence and **c)** presence of 20uM verapamil added to the bath. **B, D)** Plot of the average peak tetanic EDL, elicited by 44 subsequent 100Hz, 500ms tetanic stimulations separated by four seconds from 6-wk WT (black, n=4), ClC-1$^{-/-}$ (blue n=4), Ca$_v$1.1$^{\Delta e29}$ (orange, n=4) and Ca$_v$1.1$^{\Delta e29}$/ClC-1$^{-/-}$ red, n=4) mice in the **B)** absence and **D)** presence of 20uM verapamil added to the bath for ClC-1$^{-/-}$ (blue n=4) and Ca$_v$1.1$^{\Delta e29}$/ClC-1$^{-/-}$ red, n=4) EDLs. Dashed lines in **D)** represent average data presented in **B)** as a reference for pre-treatment. Symbols, closed circles, mean ± SEM. Statistical analysis of results in Supplemental Figure 7 are found in Supplementary Notes.

**Fig. S8**

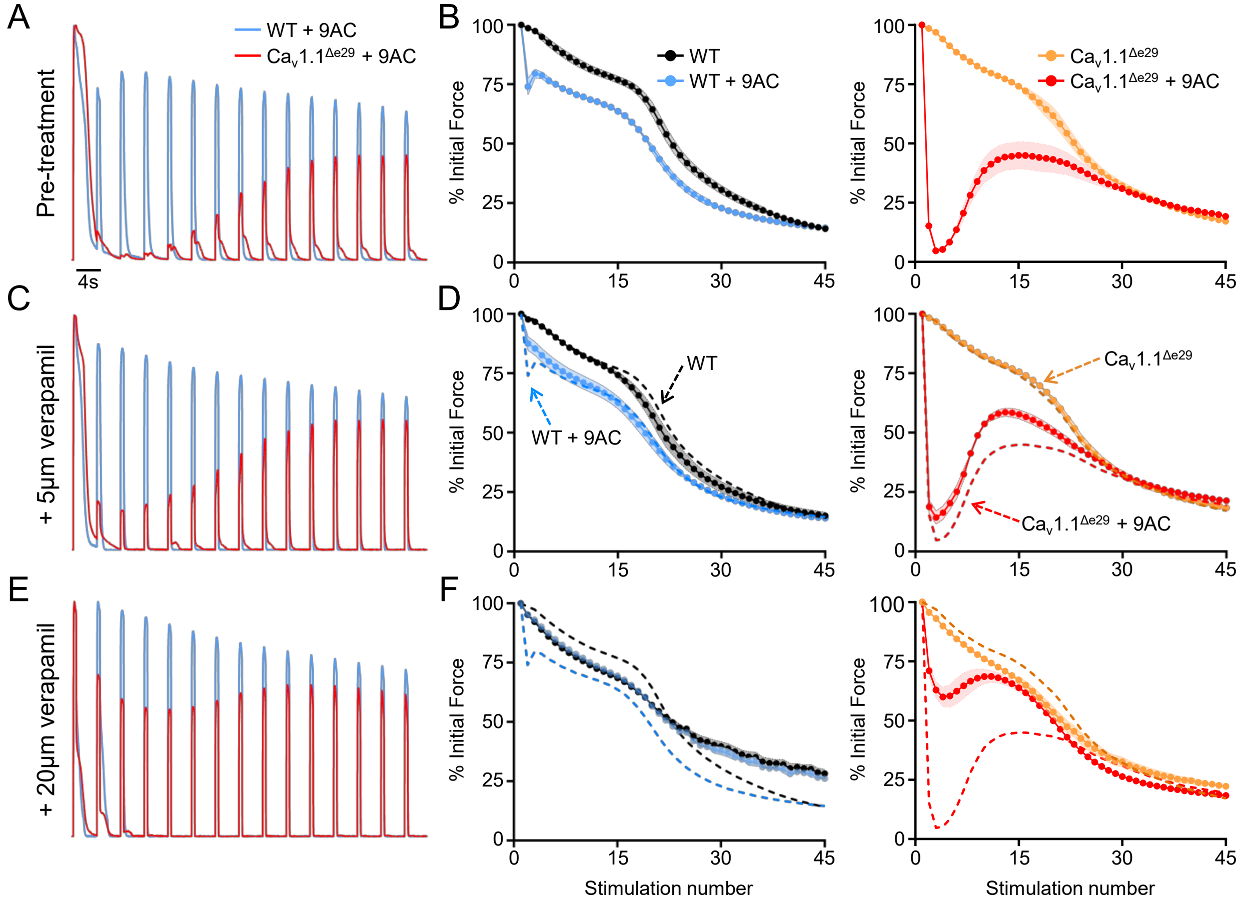


**Fig. S8. Ca$_v$1.1$^{\Delta e29}$ exacerbates transient weakness in myotonic muscle and is alleviated by verapamil.** **A)** Normalized representative force traces of the first 15 tetani (100Hz, 500ms) separated by four seconds, recorded *ex vivo* from EDLs isolated from 20-wk WT (blue) and Ca$_v$1.1$^{\Delta e29}$ (red) mice in the presence of 100mM 9-AC added to the bath (pre-treatment). **B)** Plot of the average peak tetanic EDL forces normalized to the initial stimulus, elicited by 44 subsequent 100Hz, 500ms tetanic stimulations separated by four seconds from WT (black, n=5), WT + 9-AC (blue, n=5), Ca$_v$1.1$^{\Delta e29}$ (orange, n=5) and Ca$_v$1.1$^{\Delta e29}$ + 9-AC (red, n=5) mice. **C, E)** Normalized representative force traces of the first 15 tetani (100Hz, 500ms) separated by four seconds, recorded *ex vivo* from EDLs isolated from 20-wk WT (blue) and Ca$_v$1.1$^{\Delta e29}$ (red) mice in the presence of 100mM 9-AC and **C)** 5mM verapamil or **E)** 20mM verapamil added to the bath. **D)** Plot of the average peak tetanic EDL forces normalized to the initial stimulus, elicited by 44 subsequent 100Hz, 500ms tetanic stimulations separated by four seconds from WT + 5mM verapamil (black, n=5), WT + 9-AC + 5mM verapamil (blue, n=5), Ca$_v$1.1$^{\Delta e29}$ + 5mM verapamil (orange, n=5) and Ca$_v$1.1$^{\Delta e29}$ + 9-AC + 5mM verapamil (red, n=5) EDLs. **F)** Plot of the average peak tetanic EDL forces normalized to the initial stimulus, elicited by 44 subsequent 100Hz, 500ms tetanic stimulations separated by four seconds from WT + 20mM verapamil (black, n=5), WT + 9-AC + 20mM verapamil (blue, n=5), Ca$_v$1.1$^{\Delta e29}$ + 20mM verapamil (orange, n=5) and Ca$_v$1.1$^{\Delta e29}$ + 9-AC + 20mM verapamil (red, n=5) EDLs. Dashed lines in **D, F)** represent average data presented in **B)** as a reference for pre-treatment. Symbols, closed circles, mean ± SEM. Note: Contralateral EDLs were used when possible. Statistical analysis of results in Supplemental Figure 8 are found in Supplementary Notes.

#### Fig. S9

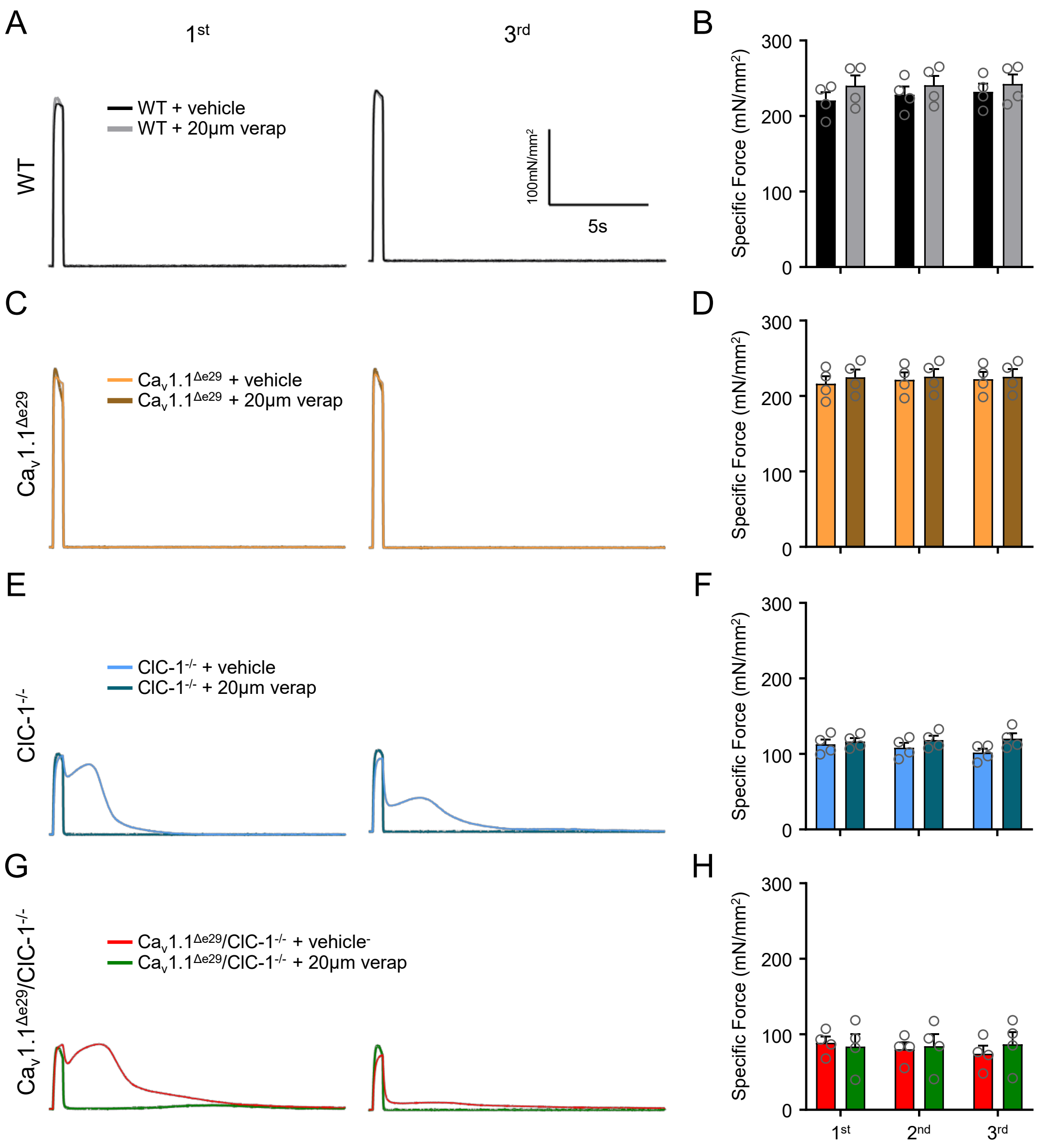


**Fig. S9. Verapamil treatment does not reduce peak contraction force of WT, $Ca_v1.1^{\Delta e29}$, ClC-$1^{-/-}$ and $Ca_v1.1^{\Delta e29}$/ClC-$1^{-/-}$ mouse muscle. A, C, E,** and **G)** Representative traces of the first (left) and third (right) tetani (150Hz, 500ms) in **A)** WT, **C)** $Ca_v1.1^{\Delta e29}$, **E)** ClC-$1^{-/-}$ and **G)** $Ca_v1.1^{\Delta e29}$/ClC-$1^{-/-}$ EDLs in the absence and presence of 20mM verapamil. Treatment depicted by colors defined in legends. **B, D, F,** and **H)** Average specific force for **B)** WT **D)** $Ca_v1.1^{\Delta e29}$ **F)** ClC-$1^{-/-}$ and **H)** $Ca_v1.1^{\Delta e29}$/ClC-$1^{-/-}$ EDLs across 3 tetanic stimulations in the absence and presence of 20mM verapamil. Treatment depicted by colors defined in legends. Symbols, open circles, individual mice; bars, mean and SEM. Statistical analysis of results in Supplemental Figure 9 are found in Supplementary Notes.

**Fig. S10**

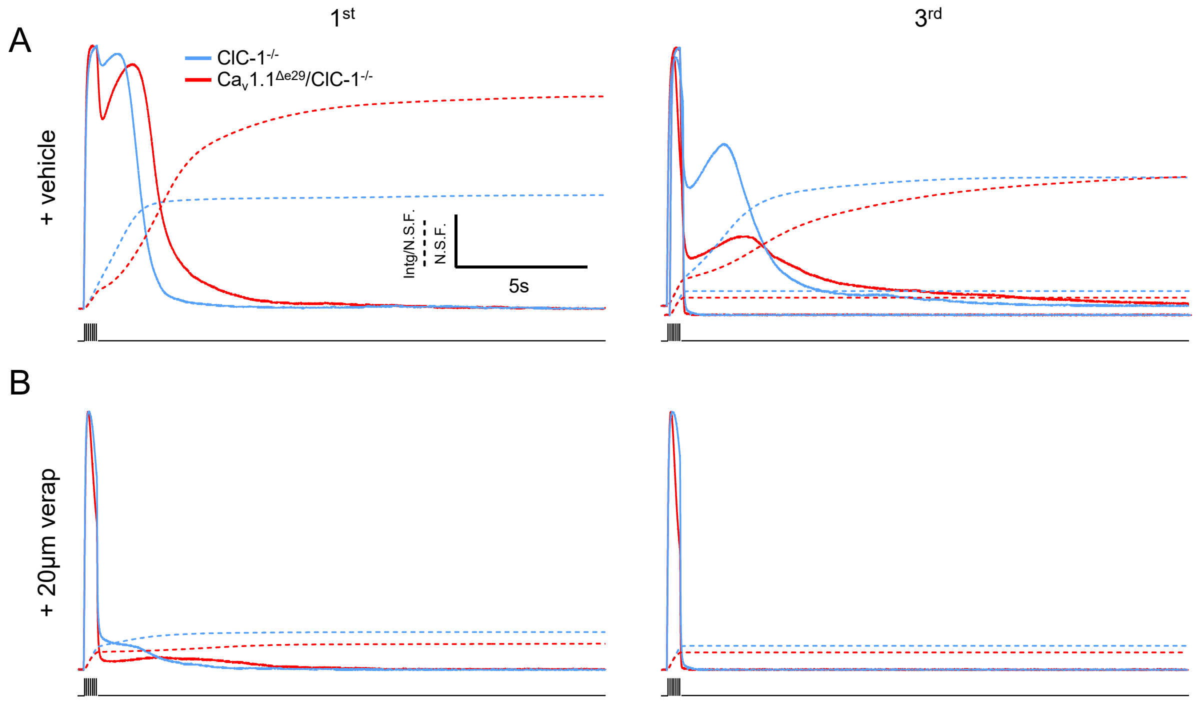


**Fig. S10. Verapamil significantly reduces myotonia in both $Ca_v1.1^{\Delta e29}$/ClC-1$^{-/-}$ and ClC-1$^{-/-}$ mouse muscle. A)** Representative trace of normalized specific force generation of the first (left) and third (right) tetani (150Hz, 500ms) in ClC-1$^{-/-}$ (blue, solid) and $Ca_v1.1^{\Delta e29}$/ClC-1$^{-/-}$ (red, sold) EDL in the absence of 20um verapamil. Dashed lines represent accumulated force. **B)** Representative trace of the first (left) and third (right) tetani (150Hz, 500ms) in ClC-1$^{-/-}$ (blue, solid) and $Ca_v1.1^{\Delta e29}$/ClC-1$^{-/-}$ (red, solid) EDL in the presence of 20 um verapamil. Dashed lines represent accumulated force. Note: Traces shown in **Fig. 3** are replotted here with expanded timescales to better observe myotonia.

**Fig. S11**

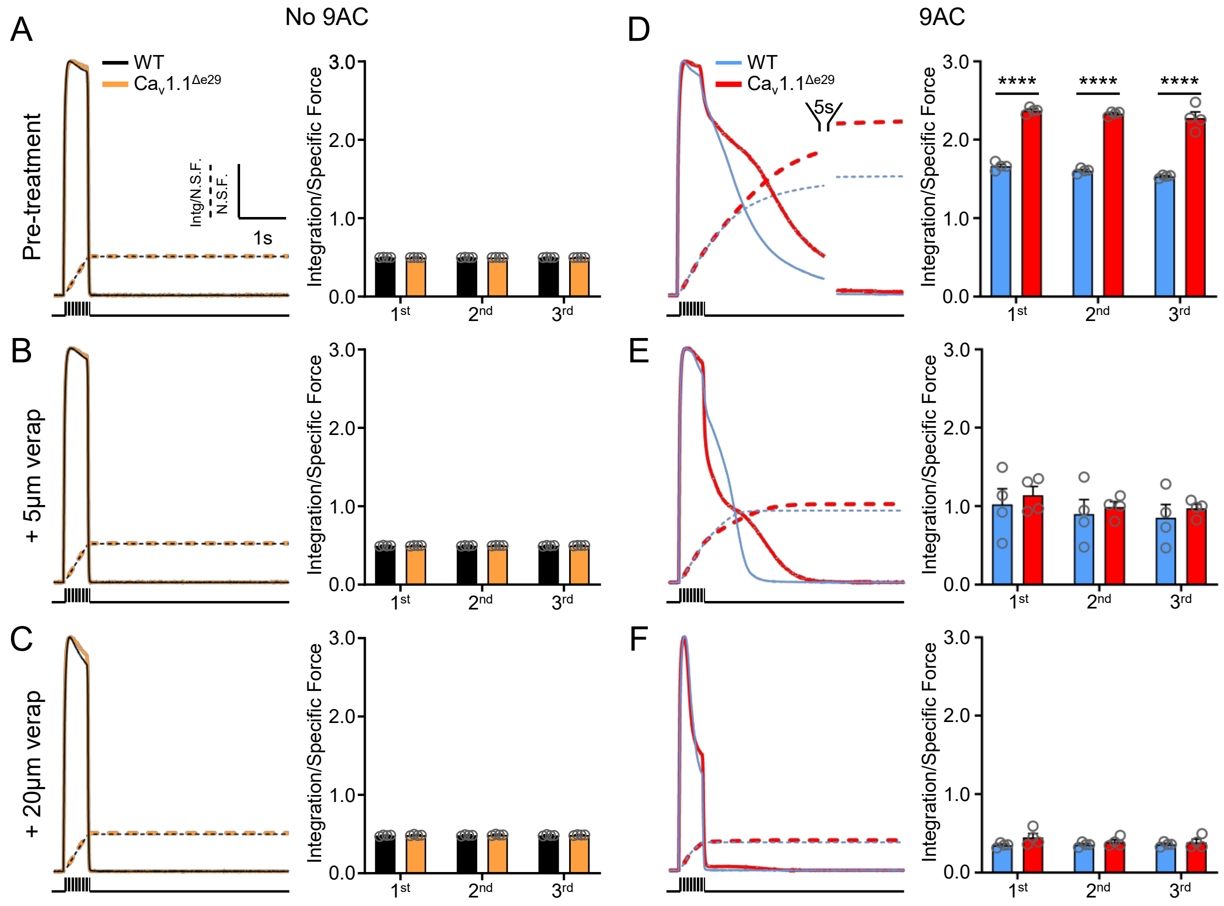


**Fig. S11. Ca$_V$1.1$^{\Delta e29}$ significantly exacerbates myotonia. A, B,** and **C, left).** Normalized representative force traces of the first of three tetani (100Hz, 500ms) separated by 3 minutes, recorded *ex vivo* from EDLs isolated from 20-wk WT (black) and Ca$_V$1.1$^{\Delta e29}$ (orange) mice in the **A, left)** absence (pre-treatment) and presence of **B, left)** 5mM and **C, left)** 20mM verapamil added to the bath (pre-treatment). Dashed lines represent accumulated force production. **A, B** and **C, right)** Plot of average integration normalized to specific force depicted in respective left panels. **D, E** and **F, left)** Normalized representative force traces of the first of three tetani (100Hz, 500ms) separated by 3 minutes, recorded *ex vivo* from EDLs incubated with 100mM 9-AC, isolated from 20-wk WT (blue) and Ca$_V$1.1$^{\Delta e29}$ (red) mice in the **D, left)** absence (pre-treatment) and presence of **E, left)** 5mM and **F, left)** 20mM verapamil added to the bath (pre-treatment). Dashed lines represent accumulated force production. **D, E** and **F, right)** Plot of average integration normalized to specific force depicted in respective left panels. Symbols, open circles, individual mice; bars, mean and SEM. Note: Contralateral EDLs were used when possible. Statistical analysis of results in Supplemental Figure 11 are found in Supplementary Notes.

**Fig. S12**

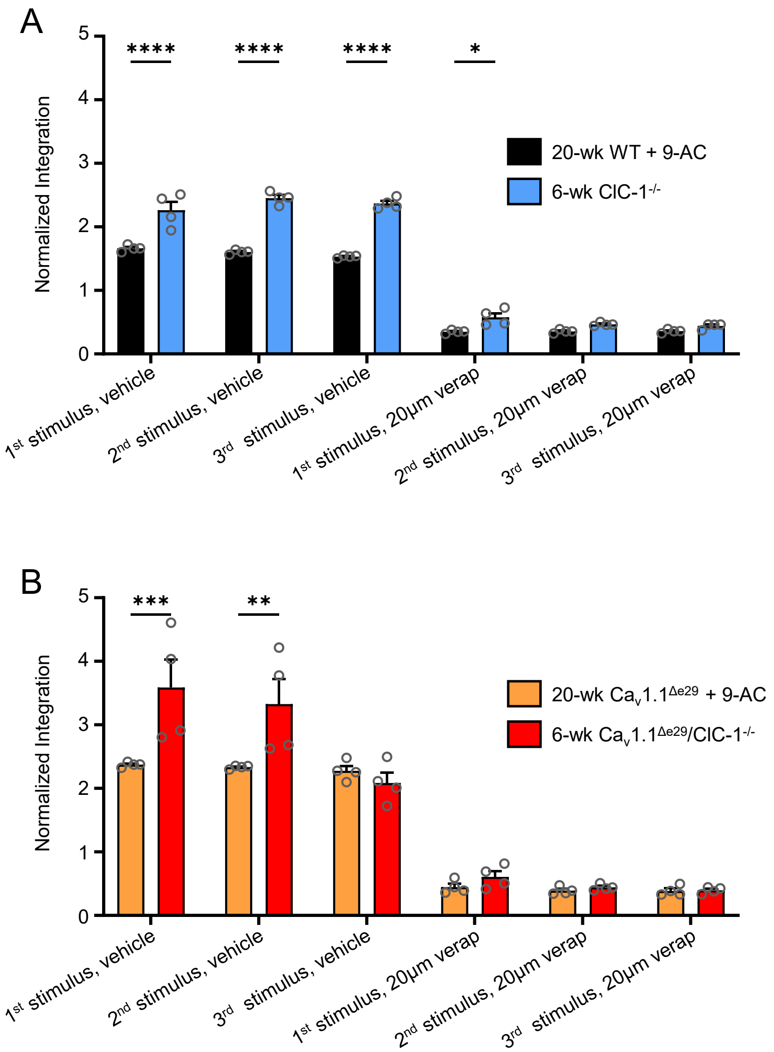


**Fig. S12. Comparison of pharmacologic and genetic myotonia. A)** Plot of average integration normalized to specific force of WT EDL + 9-AC (black) and ClC-$1^{-/-}$ (blue). **B)** Plot of average integration normalized to specific force of $Ca_V1.1^{\Delta e29}$ EDL + 9-AC (orange) and $Ca_V1.1^{\Delta e29}$/ClC-$1^{-/-}$ (red). Two-way ANOVA with multiple comparisons analysis performed, * = $P < 0.05$, ** = $P < 0.01$, *** = $P < 0.001$, and **** = $P < 0.0001$

**Fig. S13**

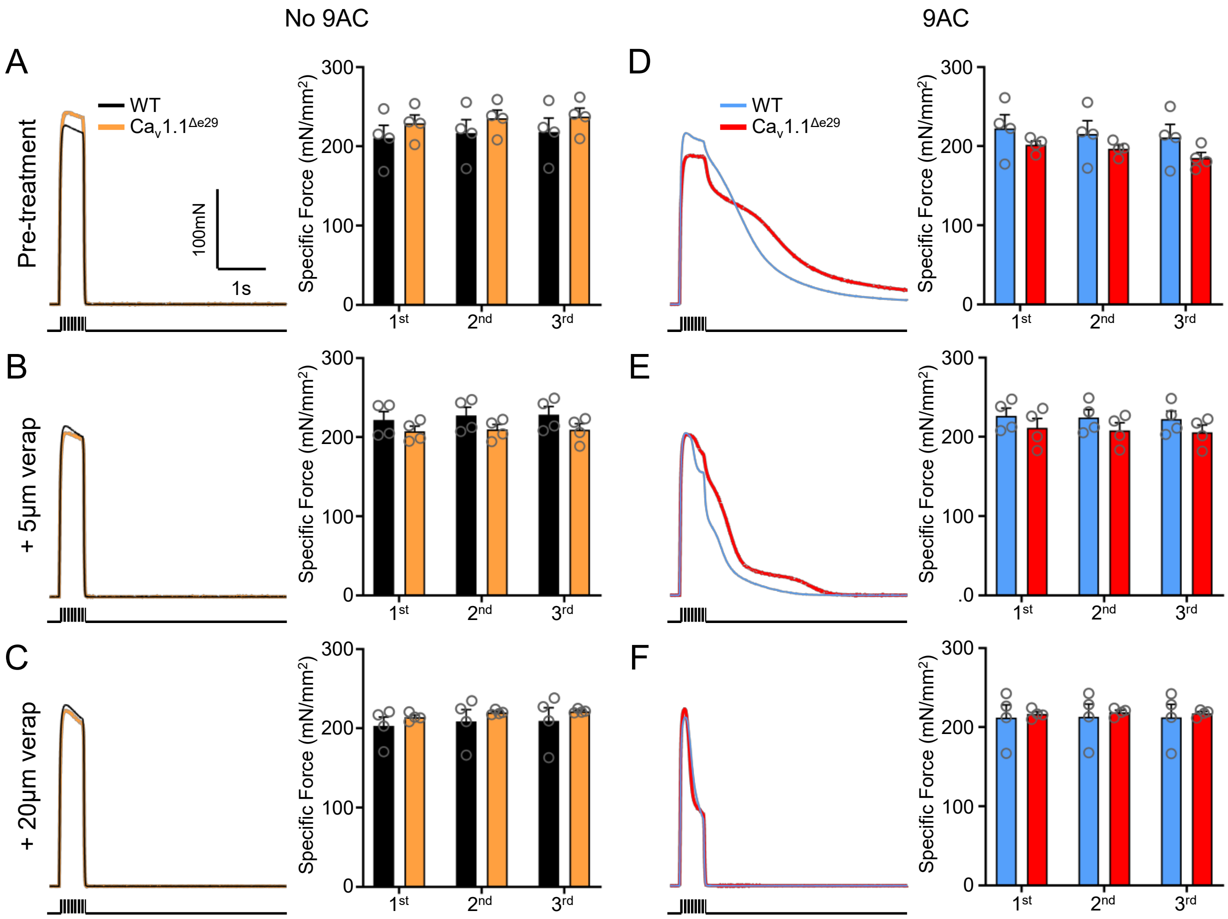


**Fig. S13. Verapamil treatment does not reduce peak contraction force of non-myotonic and myotonic WT and $Ca_v1.1^{\Delta e29}$ mouse muscle.** Representative specific force traces of the first of three tetani (100Hz, 500ms) separated by 3 minutes, recorded *ex vivo* from EDLs isolated from 20-wk WT (black) and $Ca_v1.1^{\Delta e29}$ (orange) mice in the **A, left)** absence (pre-treatment) and presence of **B, left)** 5mM and **C, left)** 20mM verapamil added to the bath (pre-treatment). Dashed lines represent accumulated force production. **A, B,** and **C, right)** Plot of average integration of specific force depicted in respective left panels. **D, E** and **F, left)** Representative specific force traces of the first of three tetani (100Hz, 500ms) separated by 3 minutes, recorded *ex vivo* from EDLs incubated with 100mM 9-AC, isolated from 20-wk WT (blue) and $Ca_v1.1^{\Delta e29}$ (red) mice in the **D, left)** absence (pre-treatment) and presence of **E, left)** 5mM and **F, left)** 20mM verapamil added to the bath (pre-treatment). Dashed lines represent accumulated force production. **D, E,** and **F, right)** Plot of average integration of specific force depicted in respective left panels. Symbols, open circles, individual mice; bars, mean and SEM. Note: Contralateral EDLs were used when possible. Statistical analysis of results in Supplemental Figure 13 are found in Supplementary Notes.

**Fig. S14**

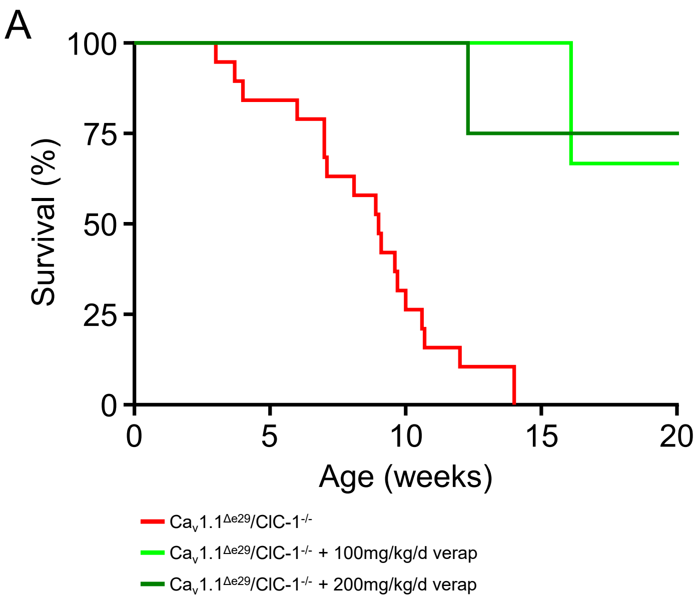


**Fig. S14. Trial of two doses of verapamil in $Ca_V1.1^{\Delta e29}/ClC\text{-}1^{-/-}$ mice results in significant rescue of survival. A)** Kaplan-Meier survival analysis of $Ca_V1.1^{\Delta e29}/ClC\text{-}1^{-/-}$ (n=19; female=9, male=10), $Ca_V1.1^{\Delta e29/\Delta e29}/ClC\text{-}1^{-/-}$ + 100mg/kg/day verapamil (n=; female=1, male=2), and $Ca_V1.1^{\Delta e29/\Delta e29}/ClC\text{-}1^{-/-}$ + 200mg/kg/day verapamil (n=4; female=2, male=2). Verapamil is dosed in mouse nutrition/hydration food cups. Statistical analysis of results in Supplemental Figure 14 are found in Supplementary Notes.

### Fig. S15

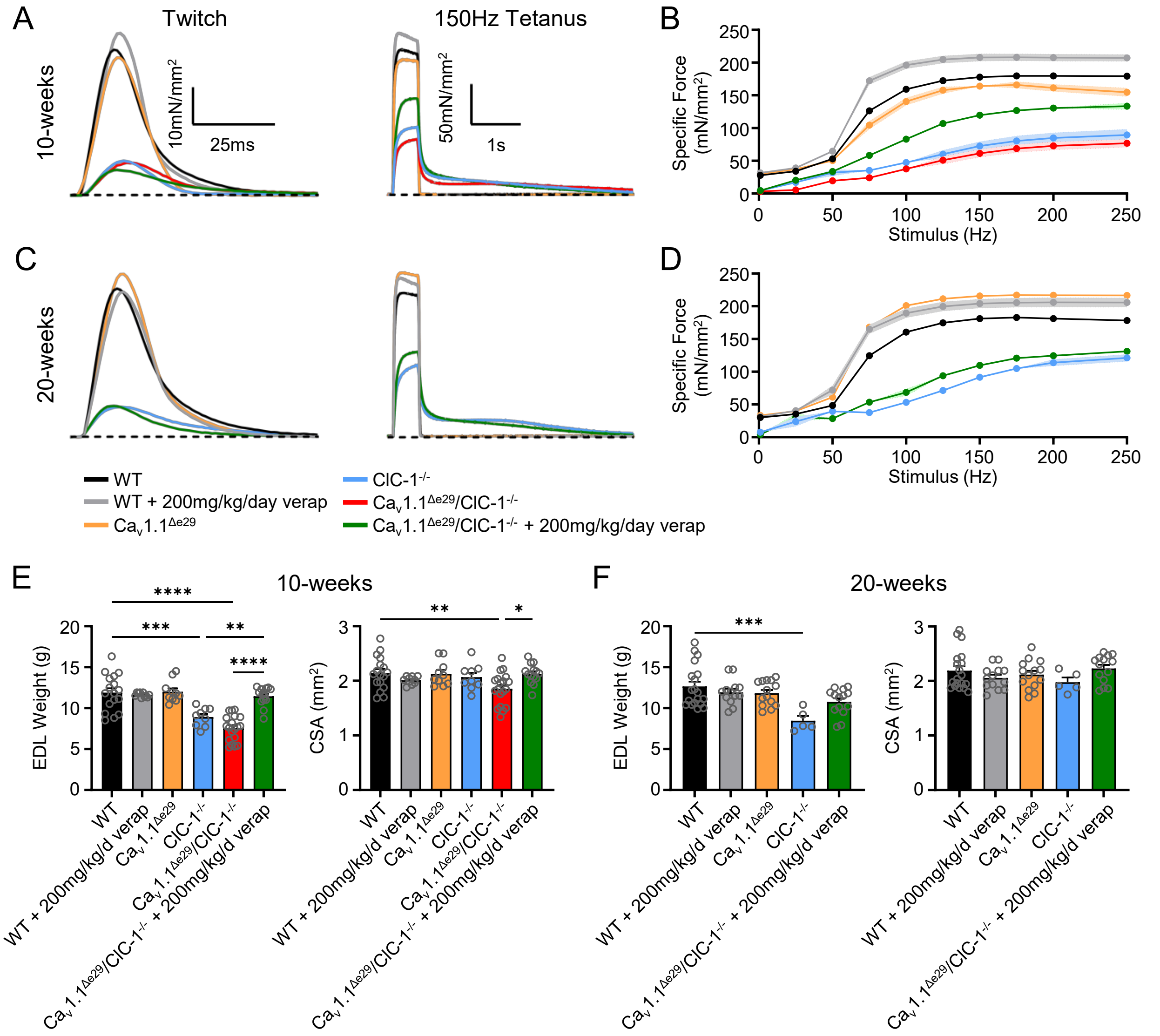


**Fig. S15. Verapamil treatment improves survival, body weight, and muscle function in $Ca_v1.1^{\Delta e29}/ClC\text{-}1^{-/-}$ mice. A, C)** Representative specific force traces elicited by twitch (left) and 150Hz (500ms) tetanic (right) stimulation of EDL muscle isolated from **A)** 10-wk and **c)** 20-wk mice. **B, D)** Plot of average stimulation frequency dependence of specific force generation from isolated EDL muscles at **D)** 10-wks and **F)** 20-wks of age in the indicated genotype and treatment groups. **E)** 10-wk and **F)** 20-wk EDL weights (left) and cross-sectional area (CSA; right). **B, E)** "n" represents individual EDLs, WT (n=17; female=8, male=9), WT + 200mg/kg/day verapamil (n=10; female=5, male=5) $Ca_v1.1^{\Delta e29}$ (n=10; female=5, male=5), $ClC\text{-}1^{-/-}$ (n=9; female=4, male=5), $Ca_v1.1^{\Delta e29}/ClC\text{-}1^{-/-}$ (n=19; female=8, male=11), and $Ca_v1.1^{\Delta e29/\Delta e29}/ClC\text{-}1^{-/-}$ + verapamil (n=14, female=7, male=7). **D, F)** "n" represents individual EDLs, WT (n=19; female=10, male=9), WT + 200mg/kg/day verapamil (n=13; female=6, male=7), $Ca_v1.1^{\Delta e29}$ (n=14; female=7, male=7), $ClC\text{-}1^{-/-}$ (n=5; female=3, male=2), $Ca_v1.1^{\Delta e29}/ClC\text{-}1^{-/-}$ + verapamil (n=14; female=7, male=7). Symbols, open circles, individual mice (**B**) or individual EDLs (**G, H**); closed circles, means ± SEM. Statistical analysis of results in Figure 15 are found in Supplementary Notes.

Table S1: sgRNA sequences for generation of force splice variant mice

| Transcript | Guide | Sequence |
|---|---|---|
| Cav1.1 | Forward | 5’- GACCTCATGTGGCCGCAGTC AGG -3’ |
| Cav1.1 | Reverse | 5’- GAGCCCCGAGAAATGGGTTG AGG -3’ |
| RyR1 | Forward | 5’ CTGGGGCTCTCTGTCGGGCTGGG 3’ |
| RyR1 | Reverse | 3’ ACAGGGGGTTTGAAAGGGTGGGG 5’ |
| SERCA1 | Forward | 5’- CCACTCCAGCTATGACTGGT GGG -3’ |
| SERCA1 | Reverse | 5’- GCGCGCGCAAGTGACCGCAG GGG -3 |

Table S2: Validation of exon 29 removal from $Ca_v1.1$ by RT-PCR

| Transcript | Direction | Exon | Sequence |
|---|---|---|---|
| Cav1.1 | Forward | 27 | 5’ CCAGTCGGAACAGATGAACCAC 3’ |
| Cav1.1 | Reverse | 31 | 5’ CCGATGACCGCGTAGATGAAGA 3’ |
| RyR1 | Forward | 66 | 5’ CCGAATCATTGTGAACAACCTGG 3’ |
| RyR1 | Reverse | 72 | 5’ GAAGGAATTCACGGACCTCCTC 3’ |
| SERCA1 | Forward | 18 | 5’ TGGGTGCAGCCACTGTAGGAG 3’ |
| SERCA1 | Reverse | 23 | 5’ AAGGGTCAGTGCCTCAGCTTTG 3’ |

Table S3: Immunohistochemistry

| Primary Antibody | Concentration | Catalogue No. | Supplier |
|---|---|---|---|
| Myosin heavy chain Type I (Isotype: MIgG2b) | 1:40 | BA-D5 | Developmental Studies Hybridoma Bank |
| Myosin heavy chain Type IIA (Isotype: MIgG1) | 1:40 | SC-71 | Developmental Studies Hybridoma Bank |
| Myosin heavy chain Type IIB (Isotype: MIgM) | 1:40 | BF-F3 | Developmental Studies Hybridoma Bank |
| Primary Antibody | Concentration | Catalogue No. | Supplier |
| AffiniPure Fab Fragment Goat Anti-Mouse IgG (H+L) | 3:100 | 115-007-003 | Jackson ImmunoResearch |
| Secondary Antibody | Concentration | Catalogue No. | Supplier |
| Goat anti-Mouse IgM (Heavy chain) Cross-Adsorbed Secondary Antibody, Alexa Fluor™ 488 | 1:1500 | A-21042 | Invitrogen |
| Goat anti-Mouse IgG1 Cross-Adsorbed Secondary Antibody, Alexa Fluor™ 568 | 1:1500 | A-21124 | Invitrogen |
| DyLight™ 405 AffiniPure Fab Fragment Goat Anti-Mouse IgG2b, Fcγ fragment specific | 1:1000 | 115-477-187 | Jackson ImmunoResearch |

Table S4: Calcium Channel Blockers Used

| Drug | Catalogue No. | Supplier |
|---|---|---|
| (±)-Verapamil hydrochloride | V4629 | Sigma-Aldrich |
